# BAF chromatin remodeling complexes inhibit immediate-early and early, but not late, transcription of herpes simplex virus 1

**DOI:** 10.1101/2025.03.27.645673

**Authors:** Sarah M Saddoris, Luis M Schang

**Author notes:** Author contributions. Sarah M Saddoris: investigation, software, visualization, writing, editing. Luis M Schang: conceptualization, funding acquisition, writing, statistical analyses, editing.

## Abstract

Herpes simplex virus 1 (HSV-1) is a highly prevalent DNA virus with a major impact on human health. The HSV-1 genome is assembled into silenced stable chromatin and minimally transcribed during latency or assembled into permissive highly dynamic chromatin and highly transcribed during lytic infections. It is unclear how HSV-1 genomes transition between chromatin states, but epigenetics, including chromatin dynamics, have been proposed to play a major role. Chromatin remodeling complexes regulate cellular chromatin dynamics and contribute to DNA transcription, replication, and repair. The BAF family of chromatin remodeling complexes includes three ubiquitously expressed complexes (cBAF, PBAF, and GBAF) and several cell type-specific ones. Some common BAF subunits interact with two HSV-1 proteins, VP16 or ICP8.

Three subunits shared by all BAF complexes and a unique subunit from each cBAF, PBAF, and GBAF were enriched in herpes nuclear domains (HND), the novel nuclear domains formed during lytic infection in which HSV-1 genomes are transcribed, replicated, and packaged. The shared ATPase SMARCA4 bound, directly or indirectly, to HSV-1 genomes. Bromodomains bind to acetylated histones and may thus be involved in this binding. However, none of four structurally unrelated inhibitors of BAF bromodomains drastically affected the recruitment of BAF subunits to HND, and neither of four commonly acetylated histone residues recognized by BAF bromodomains was enriched in the HND. BAF complexes are thus recruited to the HND by their interactions with VP16, which activates viral transcription, and ICP8. Surprisingly, the BAF complexes recruited by VP16 and ICP8 participate in inhibition of immediate-early, early, and early-late HSV-1 transcription, but not DNA replication or late transcription. We propose that BAF complexes are recruited to the HND by VP16 and ICP8, independently of their bromodomains, to inhibit viral transcription early in infection, thus contributing to the regulated cascade of gene expression. These findings also have implications to epigenetic anticancer drugs, in that it should be considered whether their use may reactivate latent herpes simplex viruses.

**Author Summary:** Herpes simplex virus 1 (HSV-1) infects over two-thirds of the world population. HSV-1 establishes latency in neurons, resulting in life-long infection. Although most infections are asymptomatic, reactivation can produce a wide range of clinical manifestations, including cold sores, stromal keratitis, and encephalitis. Available treatments do not prevent reactivation or eliminate latent viral reservoirs, as no viral proteins are expressed during latency. Epigenetic regulation plays a role during the lytic and latent cycles. Lytic HSV-1 chromatin is highly dynamic whereas latent chromatin is stable. Chromatin dynamics are regulated by multiple factors, including the chromatin remodeling complexes. Here we show that the BAF chromatin remodeling complexes regulate HSV-1 transcription during lytic infection in primary fibroblast and transformed epithelial human cells. Although these complexes are recruited to the viral genomes by viral proteins, they counterintuitively downregulate viral transcription before the onset of DNA replication. We propose that BAF complexes play a major role in the regulation of the orchestrated cascade of viral gene expression and propose to consider the potential for reactivation of herpes simplex viruses when using epigenetic inhibitors in the treatment of cancer.

## Introduction

Herpes simplex virus 1 (HSV-1) is a nuclear-replicating double-stranded DNA (dsDNA) virus that infects about two-thirds of the global population (1). During latency, HSV-1 genomes are assembled into highly stable chromatin and minimally transcribed (2, 3). During lytic infection, conversely, viral genomes are highly dynamic, accessible, and transcribed (4–7). Transcription of HSV-1 genes is highly regulated. The tegument protein VP16 complexes with HCF-1 and Oct-1 to initiate transcription of the immediate-early (IE) genes by recruiting the cellular RNA Pol II transcription complex and a number of epigenetic modifiers. The five IE proteins suppress innate antiviral responses and allow for transcription of the early (E) genes. The E proteins, including ICP8, are largely involved in metabolism and DNA synthesis. The late (L) genes are subclassified as early L (E/L) or true L. E/L gene expression increases linearly with genome copy number, while true L expression is strictly dependent on the onset of DNA synthesis. The mechanisms regulating the cascade of gene expression, including why L genes are not transcribed until the onset of DNA synthesis, are not fully understood.

Epigenetics contribute to the regulation of HSV-1 genomes during lytic and latent infection (7–15). The basic unit of chromatin is the nucleosome, which is made up of about 147 base pairs (bp) of dsDNA wrapped around an H3:H4 tetramer and two H2A:H2B dimers. Chromatin is dynamic: nucleosomes slide, breathe, assemble, and disassemble. The chromatin remodeling complexes regulate nucleosome dynamics through nucleosome sliding, ejection, and editing. They are defined by their inclusion of a SWI2/SNF2-family helicase - like protein (hereafter referred to as ATPase), which include DExx and HELICc domains to facilitate the DNA translocation required for remodeling (16).

The chromatin remodeling complexes are classified into four families based on the phylogeny of their ATPase: INO80, CHD, ISIW, and BAF (SWI/SNF). In addition to the ATPase domains responsible for DNA translocation, these complexes typically include DNA- and histone-binding domains, which facilitate recruitment and stabilization of the complexes during chromatin remodeling. These complexes are essential for cellular DNA repair, replication, and transcription, and they are mutated in over 20% of human cancers and in multiple neurodevelopmental disorders (17–23). Our understanding of their roles during viral infection, however, remains somewhat limited.

The activation domain (AD) of VP16 recruits several subunits of BAF complexes and many other chromatin remodellers to heterochromatin to induce chromatin decondensation in uninfected cells (24, 25). It also recruits a subset of them to HSV-1 genomes, but this recruitment surprisingly does not result in activation of IE transcription (26). HSV-1 gene expression, DNA replication and capsid assembly all occur within novel nuclear domains (27), which we refer to as herpes nuclear domains (HND) (8, 9). Several chromatin remodeling complex subunits are present in the HND (28, 29). The BAF ATPases SMARCA2/4 and the ISWI ATPases SMARCA1/5 interact with the HSV-1 single-stranded DNA (ssDNA) binding protein ICP8 (29). Small interfering RNA (siRNA)-mediated knockdown of SMARCA5 resulted in a twenty fold decrease in viral titers (30), whereas that of the BAF ATPases SMARCA2/BRM and SMARCA4/BRG1 did not result in inhibition of IE transcription (26); later viral functions were not evaluated. The studies of chromatin remodeling complexes during HSV-1 infection have typically focused on subunits shared by all complexes within a family, such as the ATPases, and therefore it is unclear whether the different complexes within each family play different roles during infection. These studies have also focused on global effects, whereas the regulation of the different steps in the replication cycle have not been thoroughly evaluated. We thus evaluated the three ubiquitous complexes of the BAF family: canonical BAF, cBAF; polybromo BAF, PBAF; and non-canonical GLTSCR1/1L-BAF complex, GBAF, throughout the HSV-1 lytic replication cycle.

BAF complexes are associated with transcriptional activation, splicing, DNA double stranded damage repair, and DNA decatenation following replication, among other processes (31–35). They also coordinate the inhibition of transcription at DNA damages sites to promote DNA repair (36). The specific functions of each BAF complex are not yet fully understood, but they have different preferential binding sites within the cellular genome. cBAF binds preferentially within active and primed promoters; PBAF within active promoters, gene bodies, and repressed sites; and GBAF at enhancers, promoters, and topologically associated domain (TAD) boundaries, suggesting they have different activities. Different BAF complexes can compete to activate or inhibit transcription of inducible genes (37) and PBAF is the main orchestrator of the coordination between inhibition of transcription and DNA repair (36). BAF complex structure, assembly, and function have been comprehensively reviewed (38).

Here we explore the roles of cBAF, PBAF, and GBAF during HSV-1 lytic infection in human cells. We show that different subunits of different BAF complexes are enriched in the HND. The shared ATPases SMARCA4 and SMARCA2, the shared scaffold SMARCC1, and the unique subunits ARID1A, PBRM1, and BRD9 are all enriched in the HND. SMARCA4 binds, directly or indirectly, to the viral genomes, which is consistent with BAF complexes contributing to the unique viral chromatin dynamics of HSV-1 lytic infection. BAF subunits interact with VP16 and ICP8 (24, 29), as well as with acetylated histones. However, neither of four commonly acetylated histone residues involved in recruiting BAF were enriched in the HND, and three were relatively depleted from them. None of the four structurally distinct small molecule bromodomain inhibitors drastically inhibited the recruitment of the BAF subunits either. BAF complexes are therefore recruited by their interactions with VP16 or ICP8, which activate transcription and DNA replication, respectively. As bromodomains also interact with many other proteins including transcription factors and DNA replication proteins, we tested whether the BAF complexes affected HSV-1 transcription or DNA replication. The BAF complexes inhibited the accumulation of IE, E, and E/L mRNA, but not genome copy number or true L mRNA. We propose that BAF complexes are recruited to HND early in infection through their interactions with VP16 and ICP8, independently from acetylated histones, to inhibit viral transcription before the onset of DNA replication thus supporting the strict regulation of the kinetics of expression of HSV-1 genes.

## Results

### BAF subunits are enriched in the herpes nuclear domains

To start testing the potential roles of the different BAF complexes during HSV-1 infection, we characterized the subnuclear localization of BAF subunits in HeLa and human foreskin fibroblast (HFF) cells at 6 hours post infection (hpi). We developed a novel approach to quantify nuclear signal distribution (figure 1A). Cells were stained for HSV-1 ICP4 and the protein of interest and counterstained with DAPI. ICP4 is an IE protein that localizes almost exclusively to the HND, which include the pre-replication and replication compartments. We next mask HND and nuclear non-HND regions with the ICP4 and DAPI channels, respectively, and sort the pixels from the channel of the protein of interest using the masked regions to classify each pixel as either HND, nuclear non-HND, or extranuclear. For the HND and nuclear non-HND regions, pixel intensity in each channel is distributed into twenty bins. We then normalize the frequency of each bin of pixel intensity within the HND to that within the nuclear non-HND (figure 1A).

**Figure 1.**
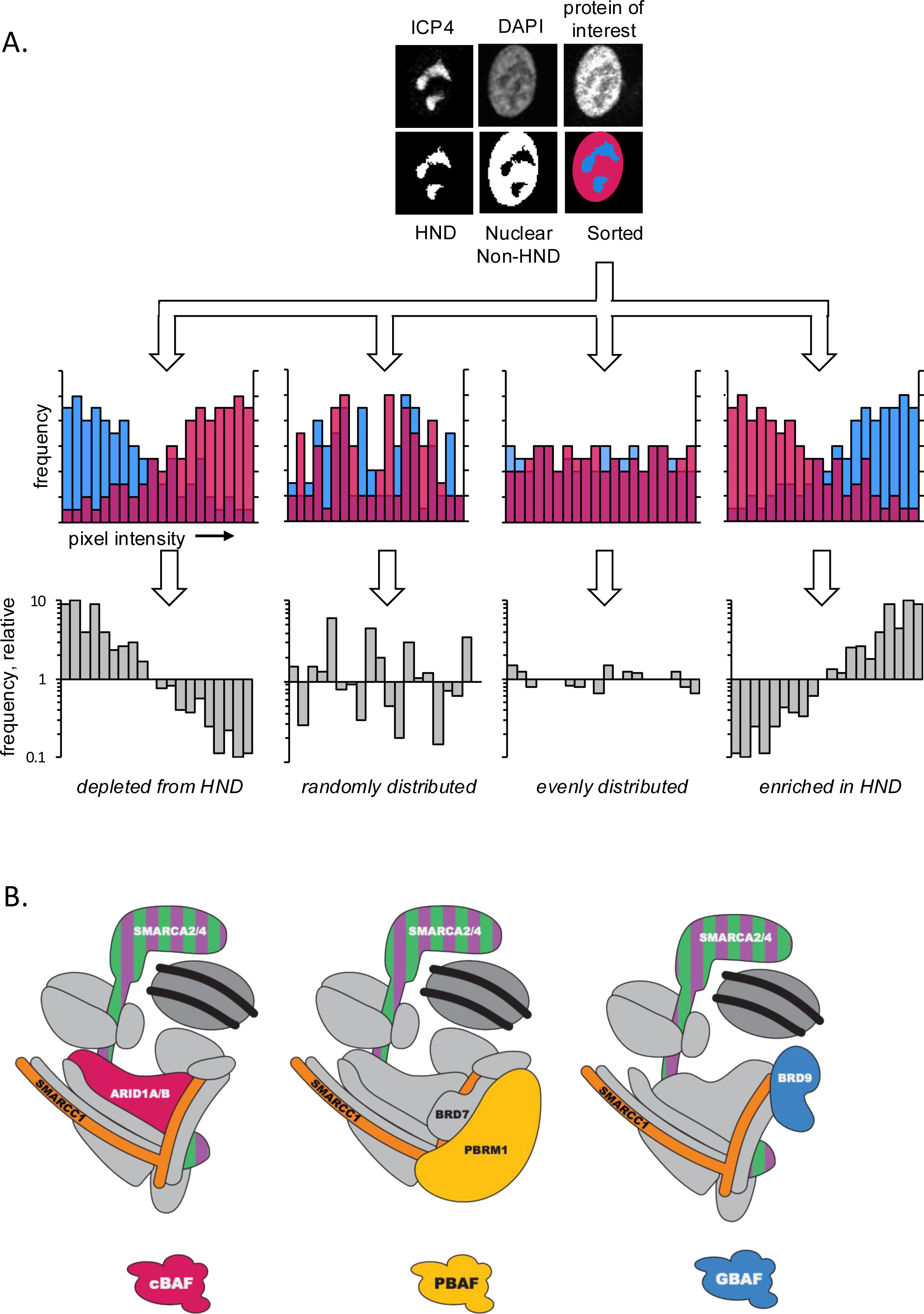
Image analysis pipeline and BAF complex structures. **A.***(i)* ICP4 and DAPI are used to mask herpes nuclear domain (HND) and nuclear, non-HND regions, respectively. *(ii)* pixels from the channel of the protein of interest are sorted as HND or nuclear non-HND (or discarded as non-nuclear). *(iii)*. For HND and nuclear non-HND regions, pixel intensities are binned, and the frequency of each bin is calculated. *(iv)* For each bin, the HND frequency is divided by the nuclear non-HND frequency. Normalized histograms are plotted. Representative histograms are shown. **B.** Cartoons depicting the different BAF complexes, modified from Varga *et al.,* (38). Subunits discussed in this manuscript are color-coded. SMARCA2/4 are the mutually exclusive ATPases. SMARCC1 is a common scaffolding subunit. ARID1A/B, PBRM1, and BRD9 are unique to cBAF, PBAF, and GBAF, respectively.

We stained for three subunits shared by all BAF complexes: SMARCA2, SMARCA4, and SMARCC1 (figure 1B, 2). SMARCA2 and SMARCA4 are the mutually exclusive ATPase subunits, and SMARCC1 is one of three scaffolding subunits required for activity (39). The ATPases SMARCA2 and SMARCA4 were similarly enriched in the HND (figure 2 A-C, Mann-Whitney, p<0.05 in all bins but 7 or 8). The direct comparison between their enrichment showed no statistically significant differences in signal intensity distribution between them (Mann-Whitney, p>0.05 for all bins). SMARCC1 was less enriched in the HND than either SMARCA2 or 4, but nonetheless the enrichment was statistically significant (Mann-Whitney, p<0.05), except in three of the highest relative intensity bins (16, 17, or 20) or bin 5 (figure 2C). Each subunit showed equivalent distribution patterns of signal intensity between HeLa and HFF cells (Figure 2C), although enrichment was always statistically significant in HFF cells (figure 2 C).

**Figure 2.**
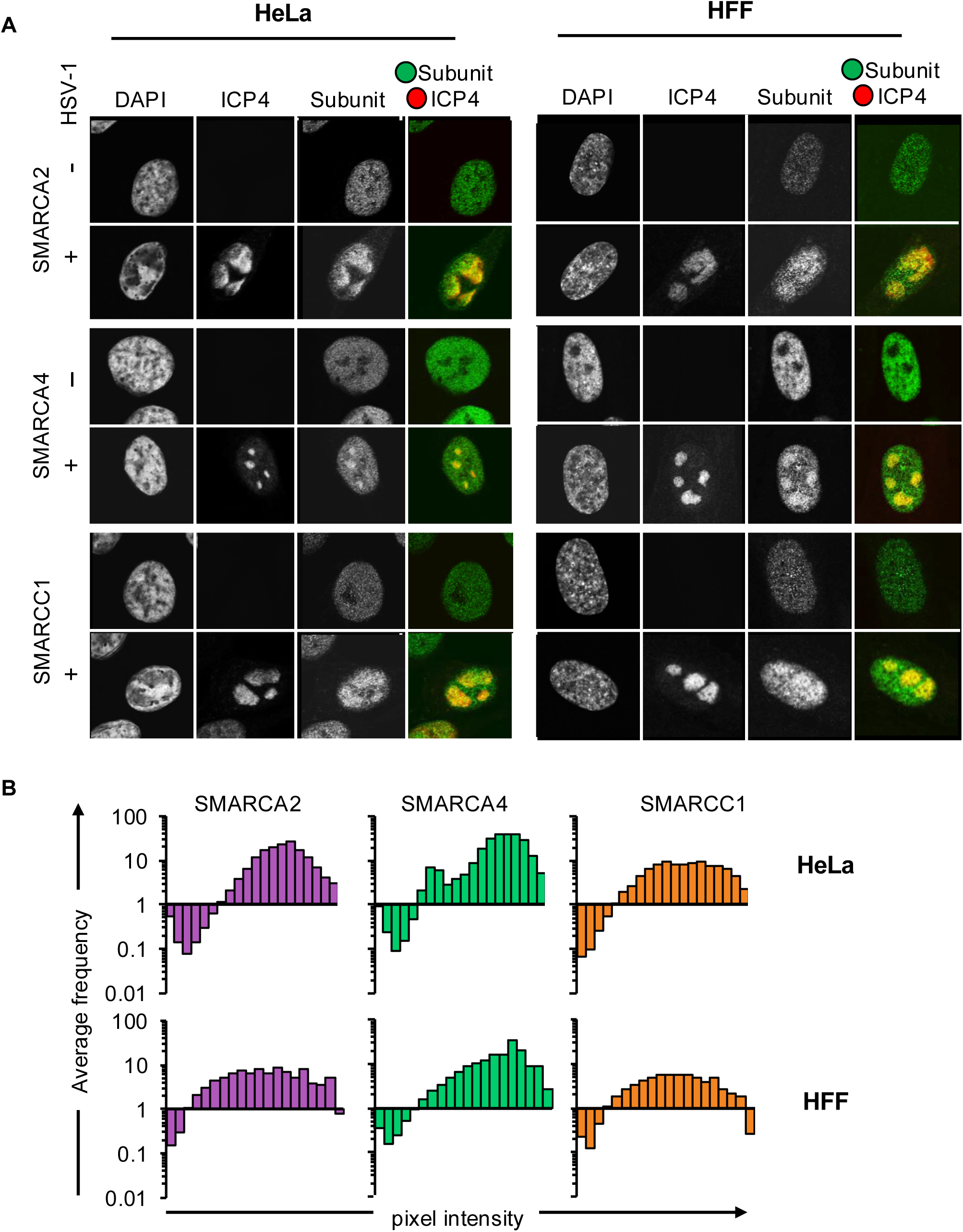
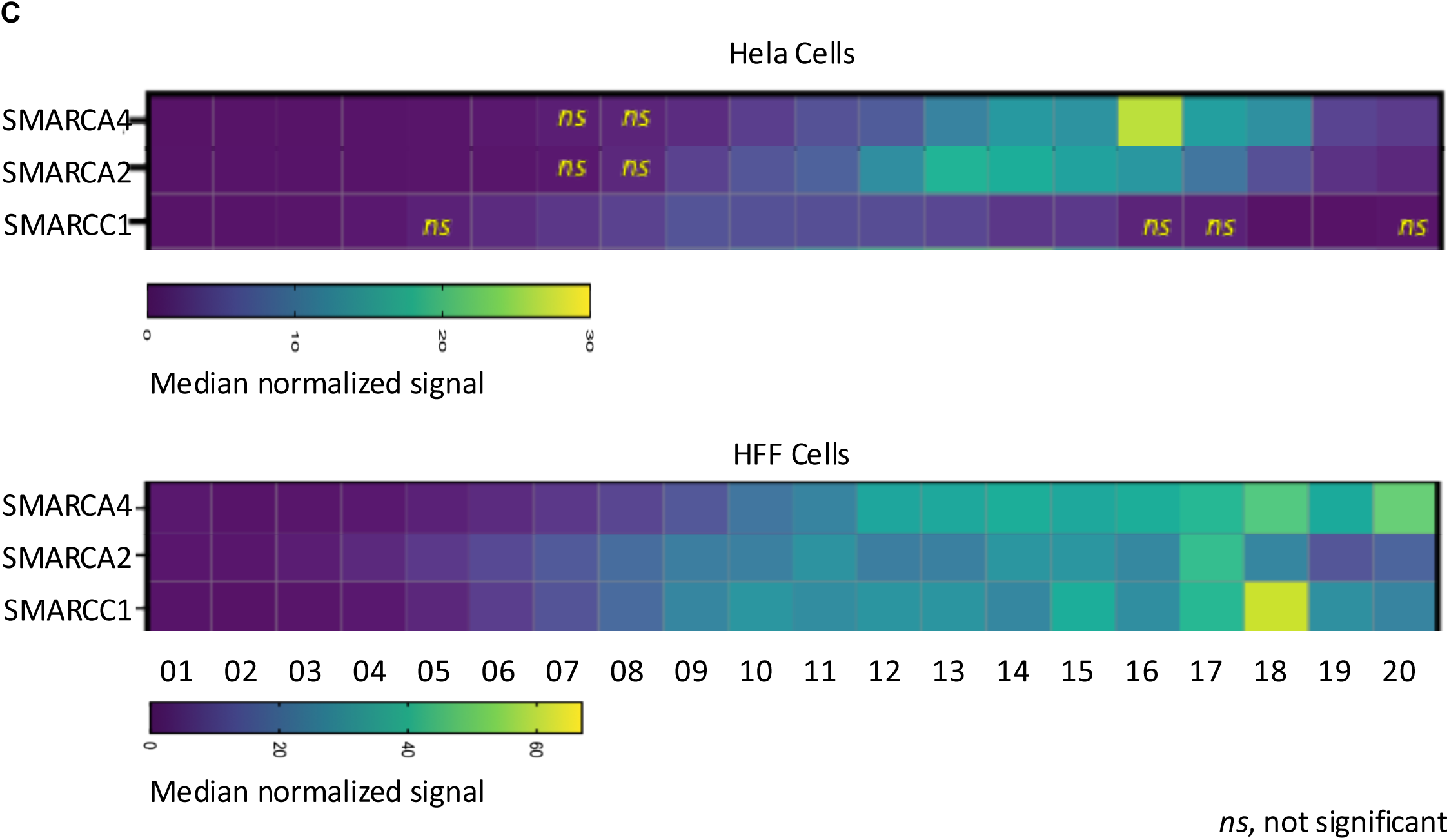
Localization of shared BAF subunits and SMARCA6 in HeLa and HFF cells. **A.**Representative micrographs of HeLa (left) and HFF (right) cells mock- or HSV-1-infected (MOI 5) fixed at 6 hpi and stained for DAPI, HSV-1 ICP4, and the subunit of interest. **B.** Analysis of the average signal distribution across three biological replicates. **C.** Heat map showing median signal intensity in the different bins (most depleted bin 1 most enriched bin 20) for each of the conserved BAF subunit in Vero or HFF cells. Frequencies in all bins except the ones indicated as *ns* were different from even distribution (Mann-Whitney, p<0.05), indicating significant depletion in the low intensity bins or enrichment in the high intensity ones.

To explore the possibility that specific BAF complexes may have different roles during HSV-1 infections, we stained for a unique subunit from each of the three ubiquitous complexes: ARID1A from cBAF, PBRM1 from PBAF, and BRD9 from GBAF (figure 1, 3). ARID1A is an essential core scaffolding subunit of cBAF (38); it does not have any bromodomains. PBRM1 includes six tandem bromodomains that modulate PBAF localization; it also binds to DNA directly. BRD9 contributes to GBAF localization and stabilization through its single bromodomain. We also attempted to evaluate a second PBAF unique subunit, BRD7, but no antibody proved specific for BRD7 by immunofluorescence (IF). The three subunits were enriched in HND in both HeLa and HFF cells (figure 3A-C). For PBRM1, the enrichment in signal distribution in HeLa cells is evidenced in the decrease in the lowest relative intensity bins (p<0.05 bins 1-3, figure 3C), while the differences in all other bins were not statistically different from an even distribution (p>0.05). PBRM enrichment in HFF cells was statistically significant in all bins, although,it was less enriched than ARID1A when directly compared to it (p<0.05 bins 2-6). ARID1A was more enriched than PBRM1 in HeLa cells, too, in the high intensity bins (p <0.05 in bins 11 to 18) and more depleted or enriched than BRD9 in the low and high intensity bins, respectively (p < 0.05 in bins 2-5 and 14-16). There were no differences in the enrichment of PBRM1 or BRD9 (p>0.05). Thus, different BAF complexes are differentially enriched in the HND, with cBAF (ARID1A) being enriched the most.

**Figure 3.**
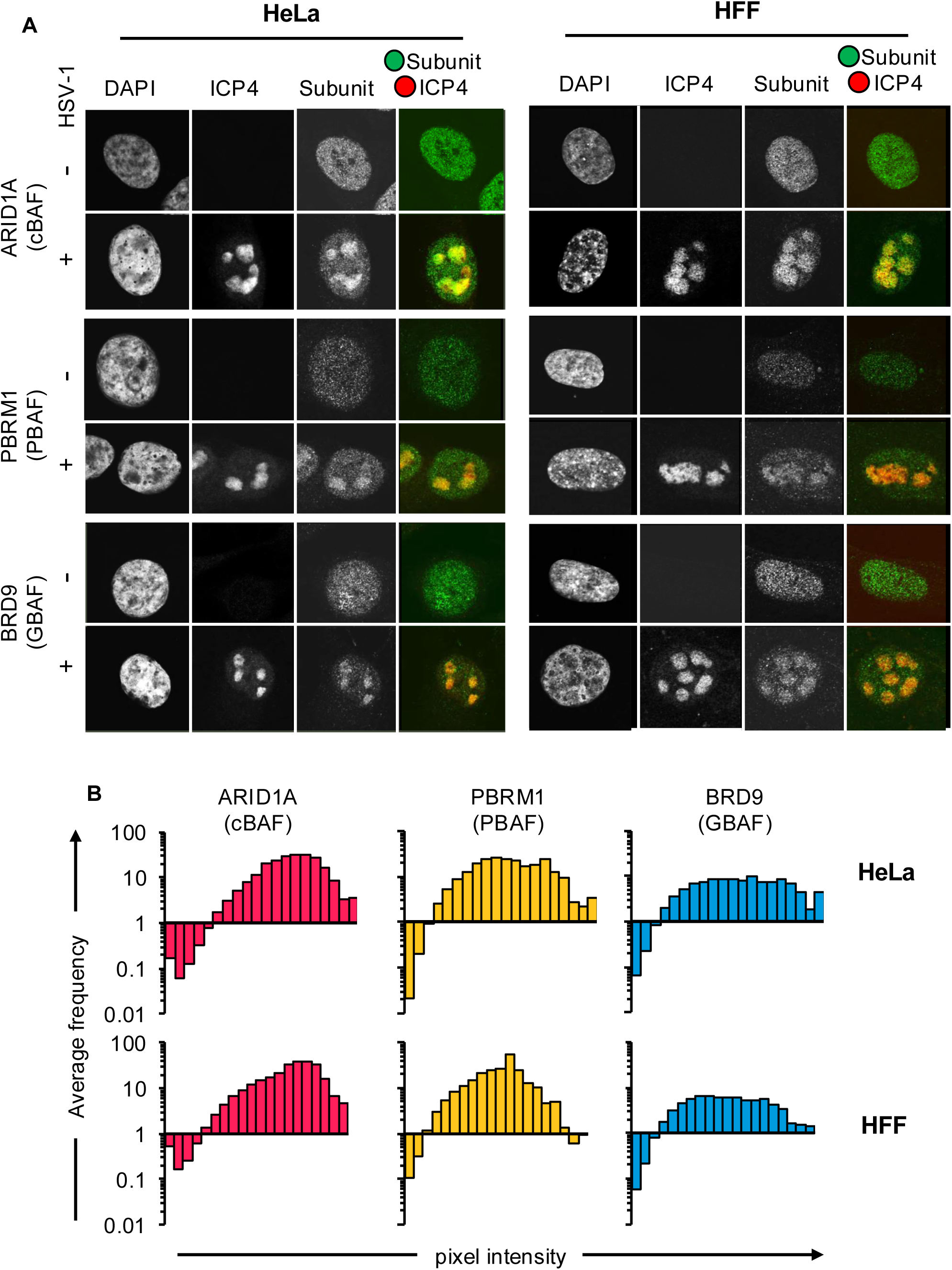
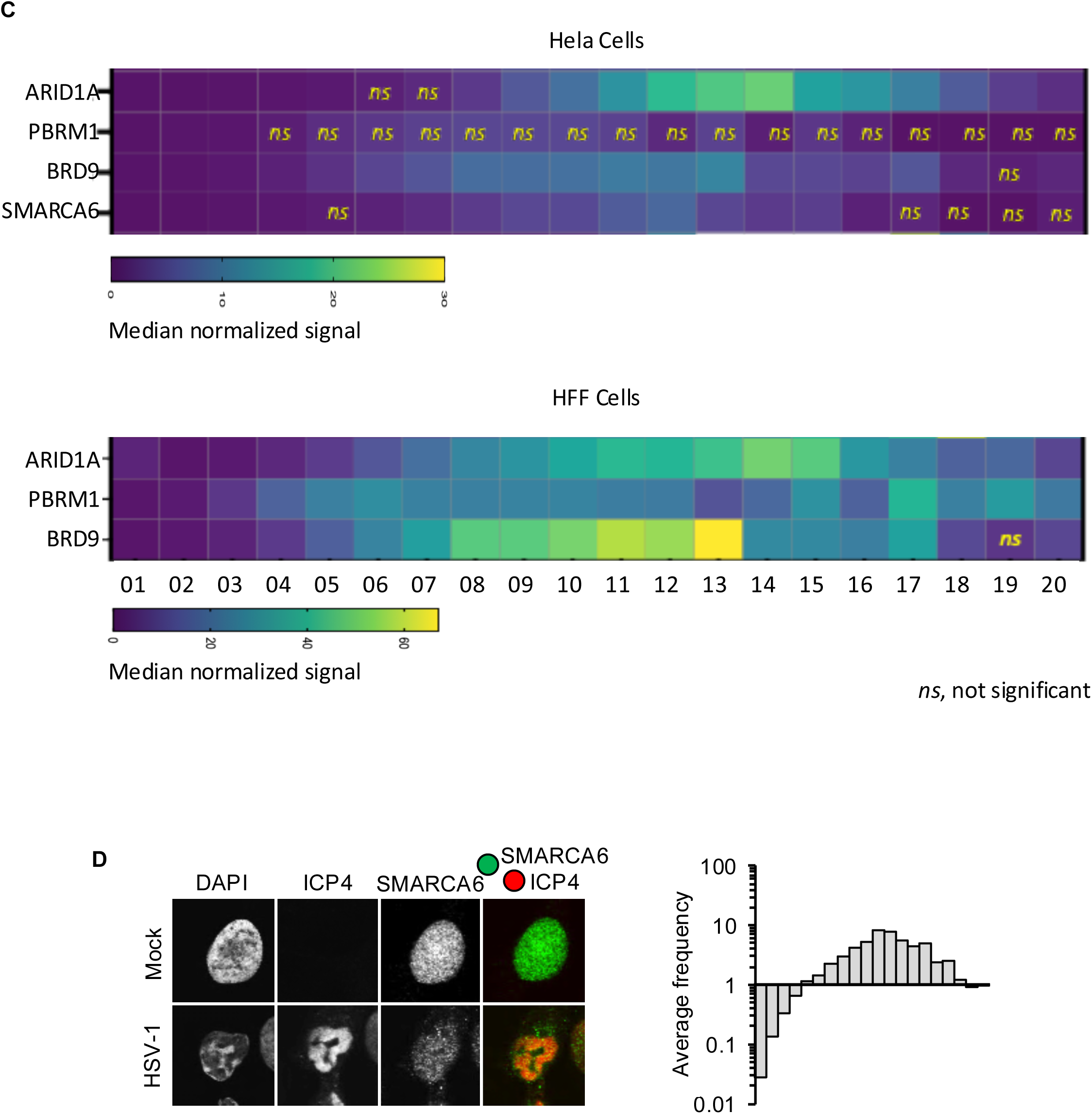
Localization of unique BAF subunits in HeLa and HFF cells. **A.**Representative micrographs of HeLa (top) and HFF (bottom) cells mock- or HSV-1-infected (MOI 5) and fixed at 6 hpi and stained for DAPI, HSV-1 ICP4, and the subunit of interest. **B.** Analysis of signal distribution across three biological replicates for each subunit and cell line. **C.** Heat map showing median signal intensity in the different bins (most depleted bin 1 most enriched bin 20) for each of the unique BAF subunits and SMARCA6 in Vero or HFF cells. Frequencies in all bins except the ones indicated as *ns* were different from even distribution (Mann-Whitney, p<0.05), indicating significant depletion in the low intensity bins or enrichment in the high intensity ones. **D.** Localization of the non BAF SMARCA6. Left, representative micrographs; right frequency distribution of the average signal intensity.

SMARCA6 is a non BAF chromatin remodeller associated with DNA methylation, and HSV-1 DNA is not methylated during lytic infection. In contrast to the BAF subunits, SMARCA6 was not enriched in the four highest intensity bins in the HND (figure 3C, D).

We used the same datasets to assess total signal intensity within HND versus non-HND regions (table 1). For each nucleus, we calculated the total pixel intensity of the protein of interest within HND and nuclear non-HND regions. We calculated the percentage of total signal intensity of the protein of interest within the HND and normalized by HND size. Although there were differences in signal distribution for individual subunits, the total average signal intensity for all BAF subunits was consistently overrepresented in HND in the two cell lines. All subunits were statistically enriched in the HND (p < 0.05), and there were no statistically significant differences in their enrichment (p>0.05), except for SMARCAC1 in comparison to SMARCA2 or 4. Consistently with its different frequency distribution, SMARCA6 was not statistically enriched in the HND.

**Table 1.** Total BAF subunit signal is over-represented in HND. IF analysis of shared and unique BAF subunits total signal within HND regions divided by total signal within nuclear, non-HND regions, normalized by HND and nuclear, non-HND sizes. >1, overrepresentation of signal in HND. n=3, average ± standard deviation; * p< 0.05; *ns, p>0.05*.

|  | HeLa | HFF |
| --- | --- | --- |
| SMARCA4 | 1.45±0.08* | 1.36±0.03 |
| SMARCA2 | 1.43±0.11* | 1.33±0.15 |
| SMARCC1 | 1.40±0.20* | 1.35±0.23 |
| PBRM1 | 1.46±0.09* | 1.28±0.04 |
| BRD9 | 1.40±0.13* | 1.27±0.01 |
| ARID1A | 1.47±0.10* | 1.33±0.10 |
| SMARCA6 | 1.17±0.09<sup>ns</sup> | ND |

### The BAF ATPase SMARCA4 binds HSV-1 genomes during lytic infection

The enrichment of the BAF subunits in HND may result from their direct or indirect binding to viral genomes. To test this possibility, we used chromatin immunoprecipitation - quantitative PCR (ChIP-qPCR). We tested SMARCA4 using a highly ChIP-validated SMARCA4 antibody (40, 41). Background immunoprecipitation with irrelevant isotype matched control antibody was subtracted before analyses. This background was typically about 10-15% of the specific signal, except for two immunoprecipitations with background levels of about 50-80%, which were considered failures and were thus not included in the analyses. We analyzed six cellular loci: IL24, LGR6, GAPDH, NPTXR a noncoding region in a gene desert in chromosome 2 (GRCh38.p14 52537120 to 52537267), and another noncoding intergenic region in chromosome 14, between SERPIN1A and SERPINA11. Consistent with previous results (40), SMARCA4 bound to about 10% of each cellular loci (12.65% average) , with the exception of the gene desert in chromosome 2 to which it bound to about 50% of input, as it would be expected (Figure 4A). We had not powered these experiments to detect any differences between cellular loci, so it is not surprising that binding to the different loci did not reach statistical significance. Binding to cellular loci was not disrupted by infection (12.65% average in mock infected cells versus 13.98% average in infected ones, or 18.76 and 18.23% if including the gene desert in chromosome 2 - figure 5A).

**Figure 4.**
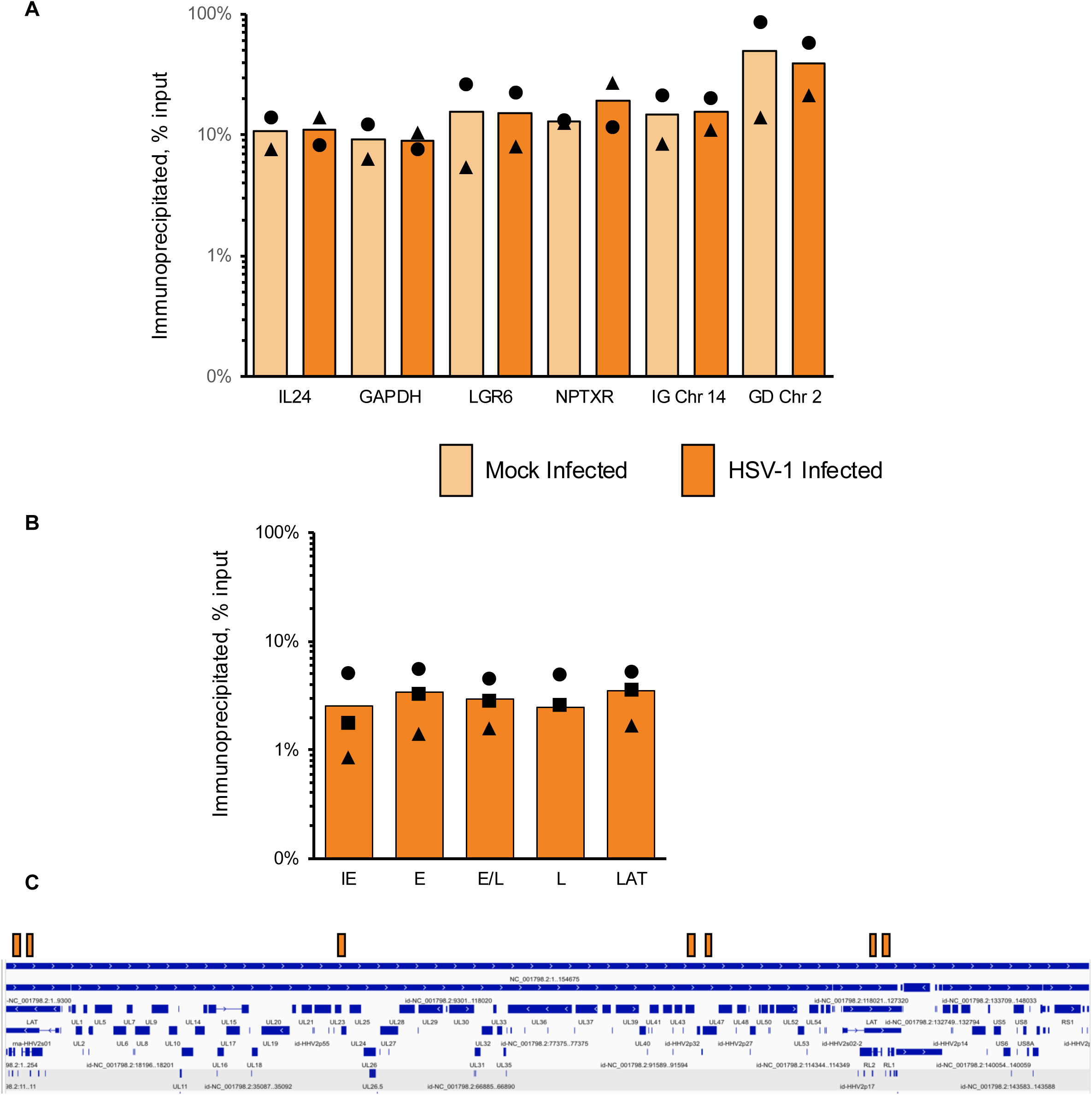
The BAF ATPase SMARCA4 binds HSV-1 genomes. HeLa cells were mock-infected or infected with HSV-1 KOS (MOI 10). Chromatin was harvested at 6 hpi and immunoprecipitated with SMARCA4 antibodies or irrelevant IgG (background). Background was subtracted from the specific IP to obtain the specific co-IP, which is plotted as percentage of input DNA before IP. SMARCA4 binding was assessed at six cellular loci **(A)** and at a locus from each kinetic class in the HSV-1 genome **(B)**. IL24, Interleukine 24, GAPDH, glyceraldehyde-3-phosphate dehydrogenase; Leucine rich repeat containing G protein-coupled receptor, NPTXR, Neuronal Pentraxin Receptor; IG Chr 14, intergenic region chromosome 14; GD Chr 2; gene desert chromosome 3; Immediate early, RL2. Early, UL23. Early late, UL46. Late, UL44. n=2 (cellular) 3 (viral). **(C)** Representation of the HSV-1 genome (blue) indicating the location of the loci tested (orange boxes).

**Figure 5.**
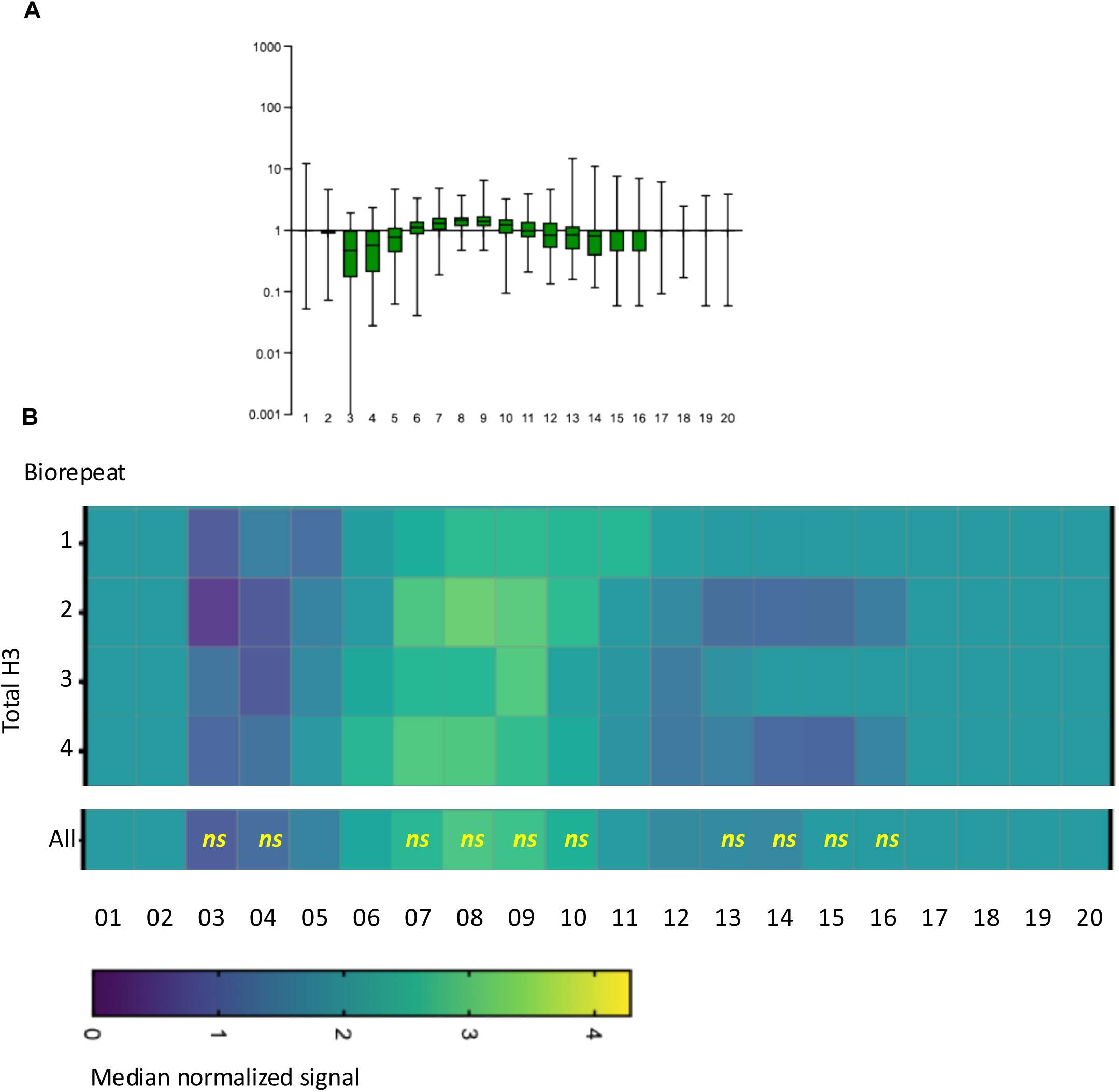
Histone H3 is not enriched neither depleted from the HND. **A.**Analysis of the average signal distribution across four biological replicates; average ± SD. **B.** Heat map showing median signal intensity in each bin in each of the 4 independent biorepeats individually and aggregated together (“all”). ns, p>0.05 (Mann-Whitney)

To test whether SMARCA4 binds HSV-1 genomes, we tested a locus from each kinetic class and distributed through the genome: RL2 (IE, short repeats), UL23 (E, unique long), UL46 (E/L, unique long), UL44 (L, unique long close to UL46), and LAT (latent transcript, repeats). SMARCA4 bound all HSV-1 loci at 6 hpi with about three-fold lower efficiency than to the cellular loci tested (figure 5B-C), with no statistical differences between viral loci (p>0.05). The difference in binding to viral and cellular loci did not reach statistical significance either but approached it at a p=0.057 (one way ANOVA), which is not unexpected since the viral chromatin is far more dynamic than the cellular one.

### Four major histone acetylation marks are depleted from HND

Bromodomains bind acetylated histones contributing to the stabilization of BAF complexes within the cellular genome (42). We tested whether histone acetylation marks were enriched in the HND, which would be consistent with the observed recruitment of BAF complexes being mediated by their bromodomains binding to the viral chromatin. We focused on four well-characterized marks: histone H3 acetylated at K4, K9, K27, or K36, normalized to total histone H3.

Histone H3 was not statistically enriched neither depleted from the HND (average 1.03±0.05, with minor enrichment in both the low and high intensity bins, Figure 5A-B). Although HSV-1 genomes are highly dynamic and are associated with acetylated histones, all tested acetylated histones except H3K4ac were relatively depleted from the HND by 15% (Table 2; p<0.05), whereas H3K4ac was neither depleted nor enriched. The signal intensity distribution also showed a general depletion for K9ac, K27ac, and K36ac , which were all relatively enriched in the low intensity bins and depleted form the mid and high intensity ones (figure 6, p<0.05), whereas H3K4ac was only depleted from the high intensity bins. The BAF complexes are thus not preferentially recruited to the HND by their interactions with acetylation of the tested histone residues.

**Figure 6.**
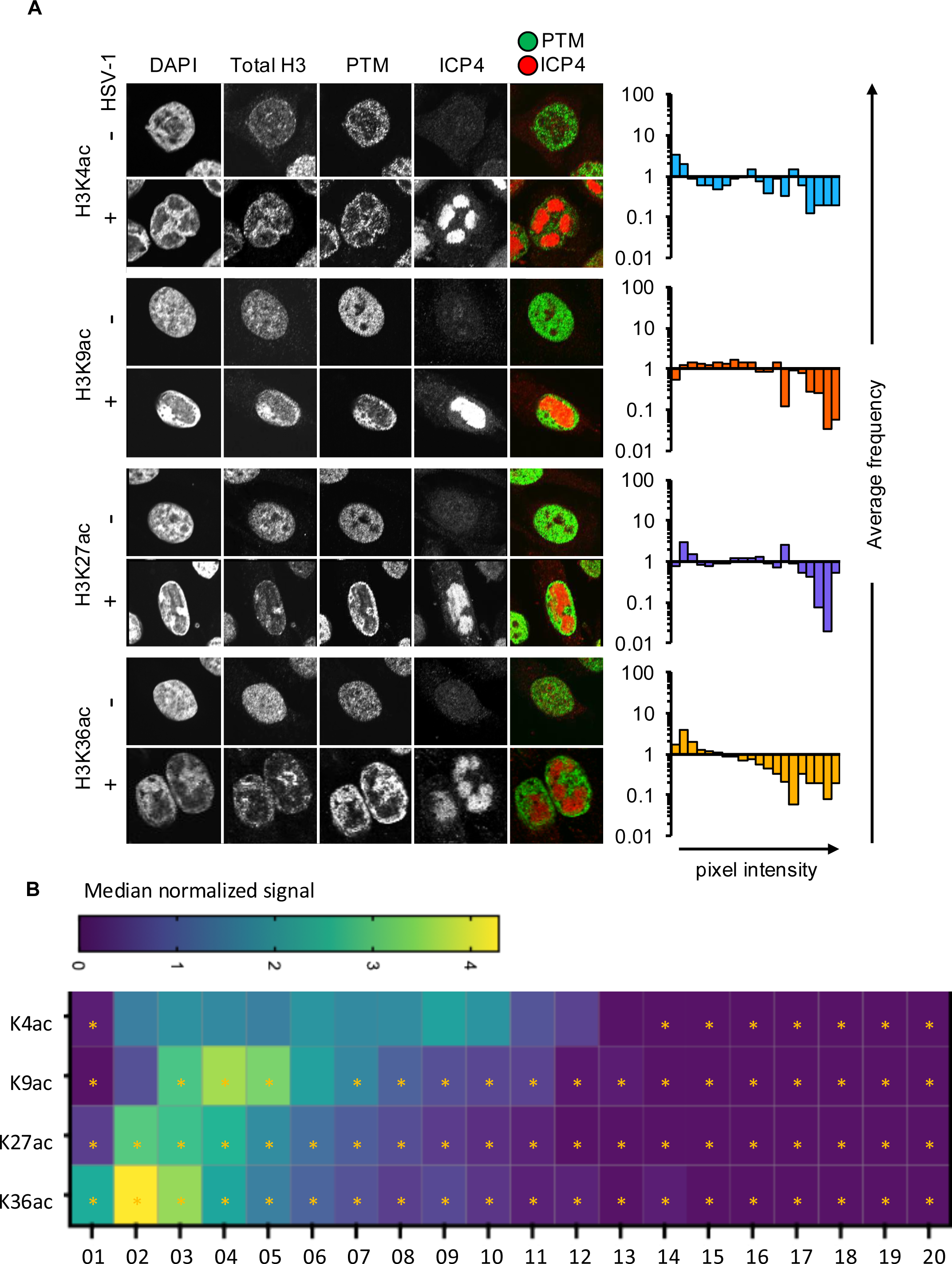
Four major histone acetylation marks are relatively depleted from HND. Localization and signal distribution of histone H3 acetylated at K 4, 9, 27, or 36 during HSV-1 infection. **A.** Representative micrographs of HeLa cells mock- or HSV-1-infected, fixed at 6 hpi, and stained for DAPI, HSV-1 ICP4, H3, and the H3 post-translational modification (PTM) of interest. Analysis of the average signal distribution across three biological replicates is shown to the right for each PTM. **B.** Heat map presenting median signal intensity for the frequency distribution of the normalized signal for each histone modification in each bin. *, statistically different from even distribution, indicating depletion or enrichment in the respective bin (Mann-Whitney, p<0.05)

**Table 2.** Total histone H3 or H3 acetylated at K4 are neither enriched nor depleted from HND, whereas H3 acetylation at K9, 27 , or 36 are relatively depleted from them . IF analysis of total or acetylated histone H3 total signal within HND regions divided by total signal within nuclear non-HND regions, normalized by HND and nuclear non-HND sizes. <1, signal underrepresented in HND. n=2, average ± standard deviation; * p< 0.05; *ns, p>0.05*.

|  |  |
| --- | --- |
| Total H3 | 1.03±0.05 *ns* |
| H3K4ac | 1.03±0.17 *ns* |
| H3K9ac | 0.85±0.06* |
| H3K27ac | 0.84±0.07* |
| H3K36ac | 0.86±0.07* |

### Small molecule bromodomain inhibitors do not prevent BAF subunit recruitment into HND

To further test whether the bromodomains were involved in the recruitment of BAF subunits to the HND, we took advantage of the multiple small molecule inhibitors of BAF bromodomains that have been developed as potential anticancer drugs (43). We selected four inhibitors from the Structural Genomics Consortium: LP99, PFI-3, BI-9564, and I-BRD9 (figure 7A). PFI-3 targets the bromodomains of SMARCA2, SMARCA4, and PBRM1, a unique subunit of PBAF. BI-9564, LP99, and I-BRD9 target the bromodomains of BRD7 and BRD9, which are unique subunits of PBAF and GBAF, respectively (44–46) (figure 7B). Cytotoxicity was assessed in HeLa and HFF cells at 18 h post treatment, the longest time used for any evaluation of any viral function but DNA replication (table 3). LP99 (10 μM), PFI-3 (32 μM), and BI-9564 (15 μM) had no major obvious effects on cell viability, while I-BRD9 (10 μM) resulted in about 25% decrease in cell viability (table 3), in agreement with their known potencies in cell culture.

**Figure 7.**
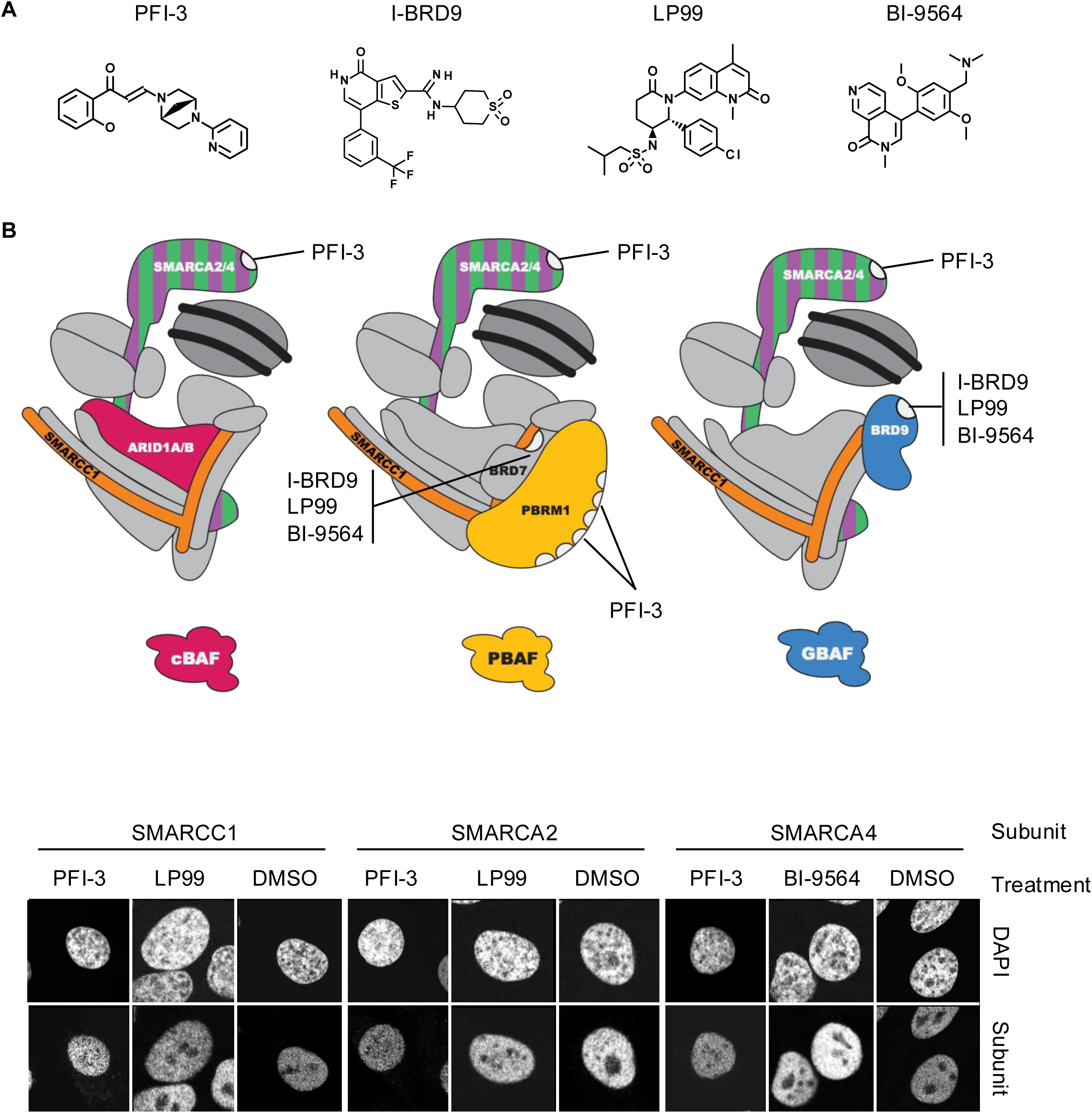
Small molecule bromodomain inhibitors targeting different BAF complexes have no effects on the distribution of the complexes in mock infected cells. **A.**Structures of the small molecule bromodomain inhibitors selected from the Structural Genomics Consortium. **B.** BAF complex structures modified from Varga et al. Subunits discussed in this manuscript are colored. Relevant bromodomains are shown in white, and the inhibitors that target each of them are listed. **C.** HeLa cells were mock-infected, treated with DMSO or a small molecule inhibitor, fixed at 6 hpi, and stained for DAPI and the subunit of interest.

**Table 3.** Small molecule bromodomain inhibitors are well-tolerated by HeLa and HFF cells. Cytotoxicity of small molecule bromodomain inhibitors was assessed by measuring ATP levels in HeLa and HFF cells treated with DMSO vehicle, LP99 (10 μM), PFI-3 (32 μM), BI-9564 (15 μM), or I-BRD9 (10 μM). Cell division rate relative to DMSO treated cells is shown. n=3, average ± standard deviation

| Compound | [μM] | HeLa | HFF |
| --- | --- | --- | --- |
| LP99 | 10 | 0.92 ± 0.12 | 1.01 ± 0.10 |
| PFI-3 | 32 | 1.00 ± 0.14 | 1.01 ± 0.05 |
| BI-9564 | 15 | 0.98 ± 0.02 | 1.05 ± 0.09 |
| I-BRD9 | 10 | 0.74 ± 0.23* | 0.72 ± 0.03 |

The small molecule inhibitors did not disrupt the global nuclear distribution of the shared BAF subunits in mock-infected HeLa cells, as expected (figure 7C). There was no global depletion of BAF subunit enrichment in the HND in cells infected and treated with any of the inhibitors in comparison to those treated with DMSO vehicle either (p >0.05, tables 4, 5). The analysis of the signal distribution, however, revealed that the inhibitors did affect somewhat the distribution of most BAF subunits (figure 8-10). PFI-3 and BI-9564 induced a decrease in signal in the low intensity bins (p <0.05) for all subunits except PBRM1, whose signal was already decreased without any inhibitor, while LP99 induced a decrease in the low intensity bins for only SMARCC1 and BRD9 (figure 10, p <0.05). These results are consistent with the status of histone acetylation in the HND in that they indicate that the BAF complexes are recruited mostly independently of interactions between their bromodomains and acetylated histones, although targeting the bromodomains in SMARCA2 and 4, and a subset of those in PBRM1, with PFI-3, or that in BRD9 with LP99, marginally affects the distribution of most subunits. These results, together with the interactions of the BAF subunits with VP16 or ICP8 (24, 29), indicate that the BAF complexes are recruited into the HND by these viral proteins.

**Figure 8.**
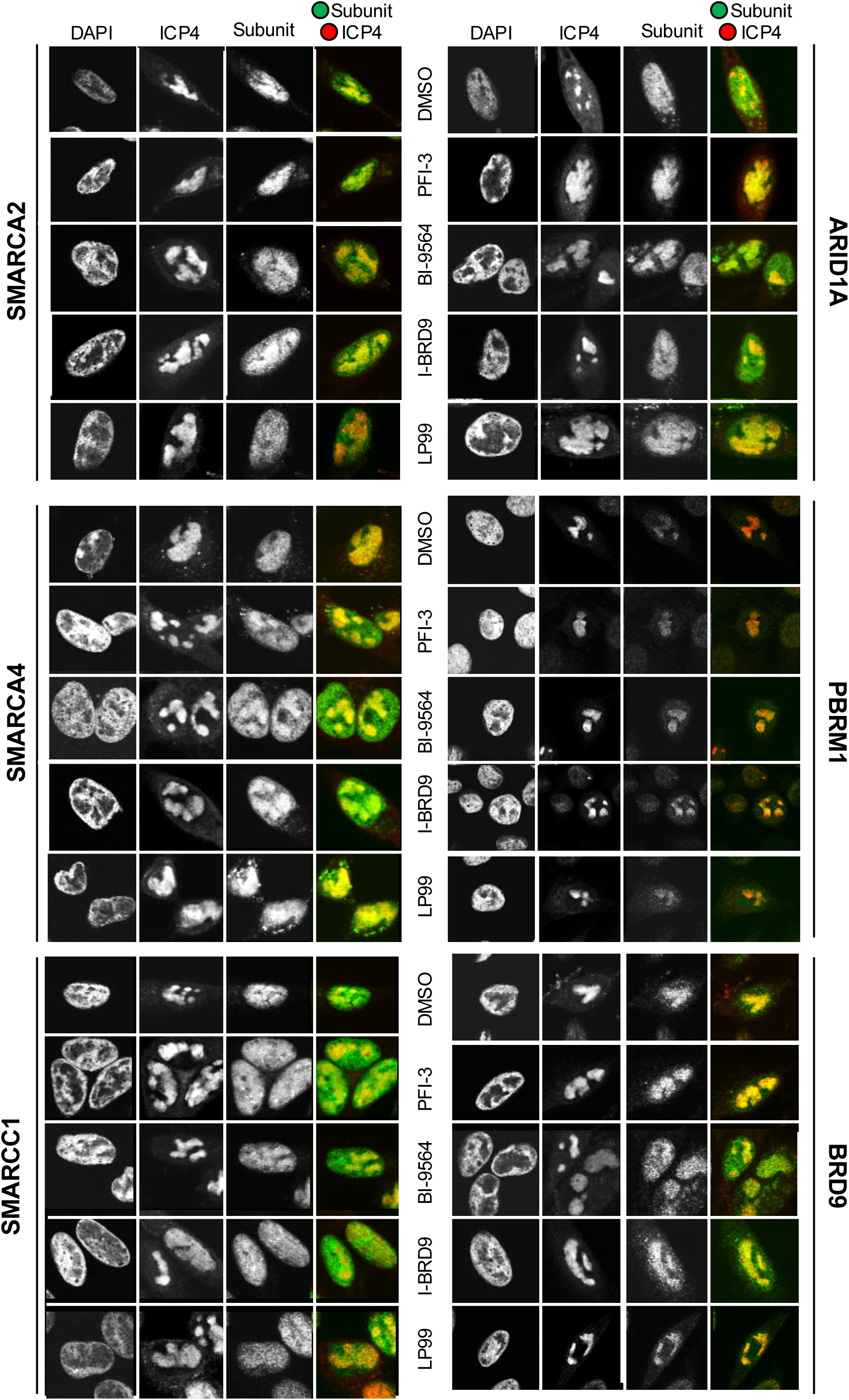
Distribution of shared BAF subunits during HSV-1 infection in the presence of small molecule bromodomain inhibitors. **A.**Representative micrographs of HeLa cells infected with HSV-1, treated with DMSO vehicle, LP99 (10 μM), PFI-3 (32 μM), BI-9564 (15 μM), or I-BRD9 (10 μM), fixed at 6 hpi, and stained for DAPI, HSV-1 ICP4, and the subunit of interest; n=2. **B.** Histograms presenting the average signal distribution of the shared BAF subunits in the presence of each inhibitor; n=2.

**Figure 9.**
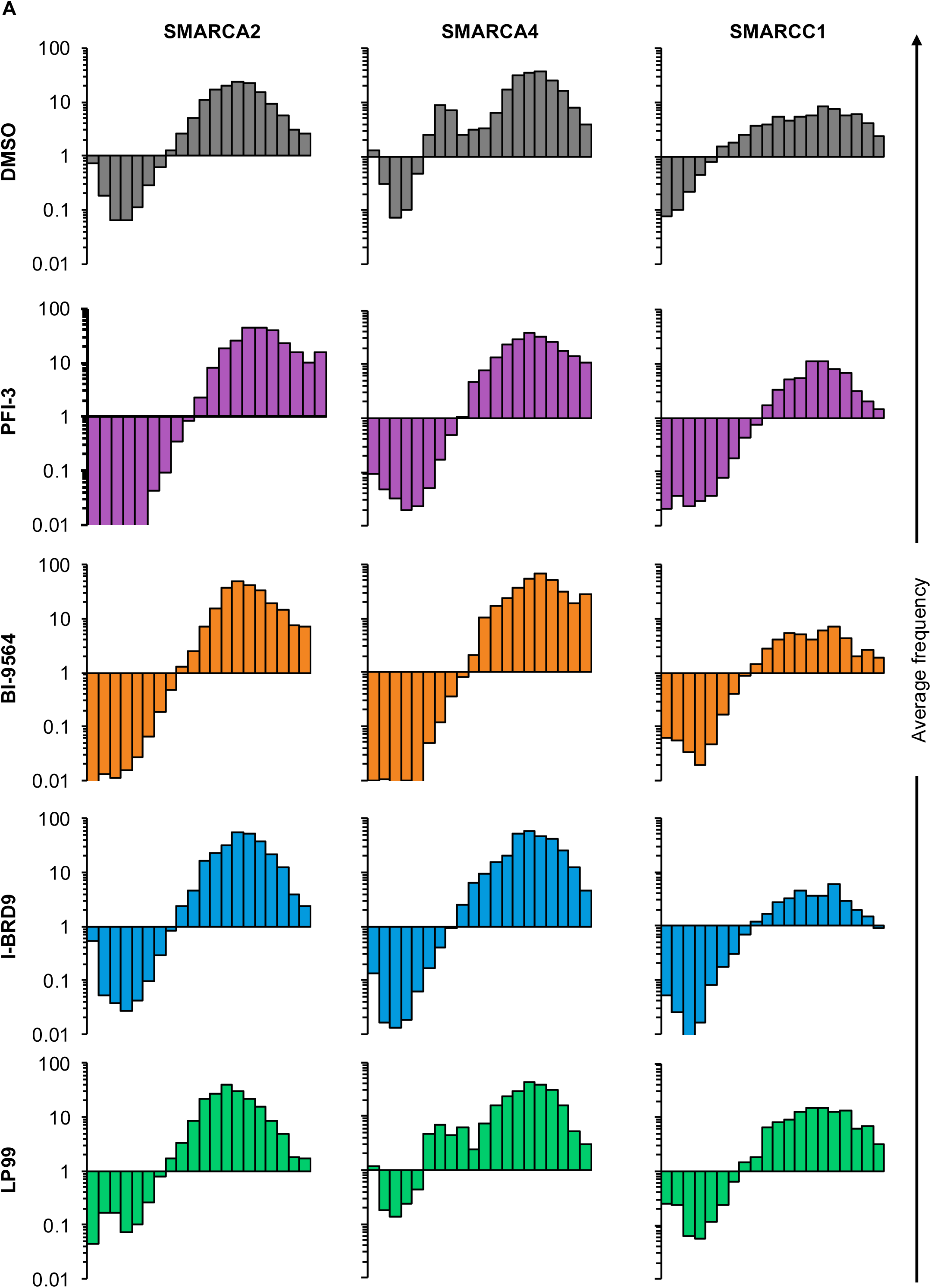
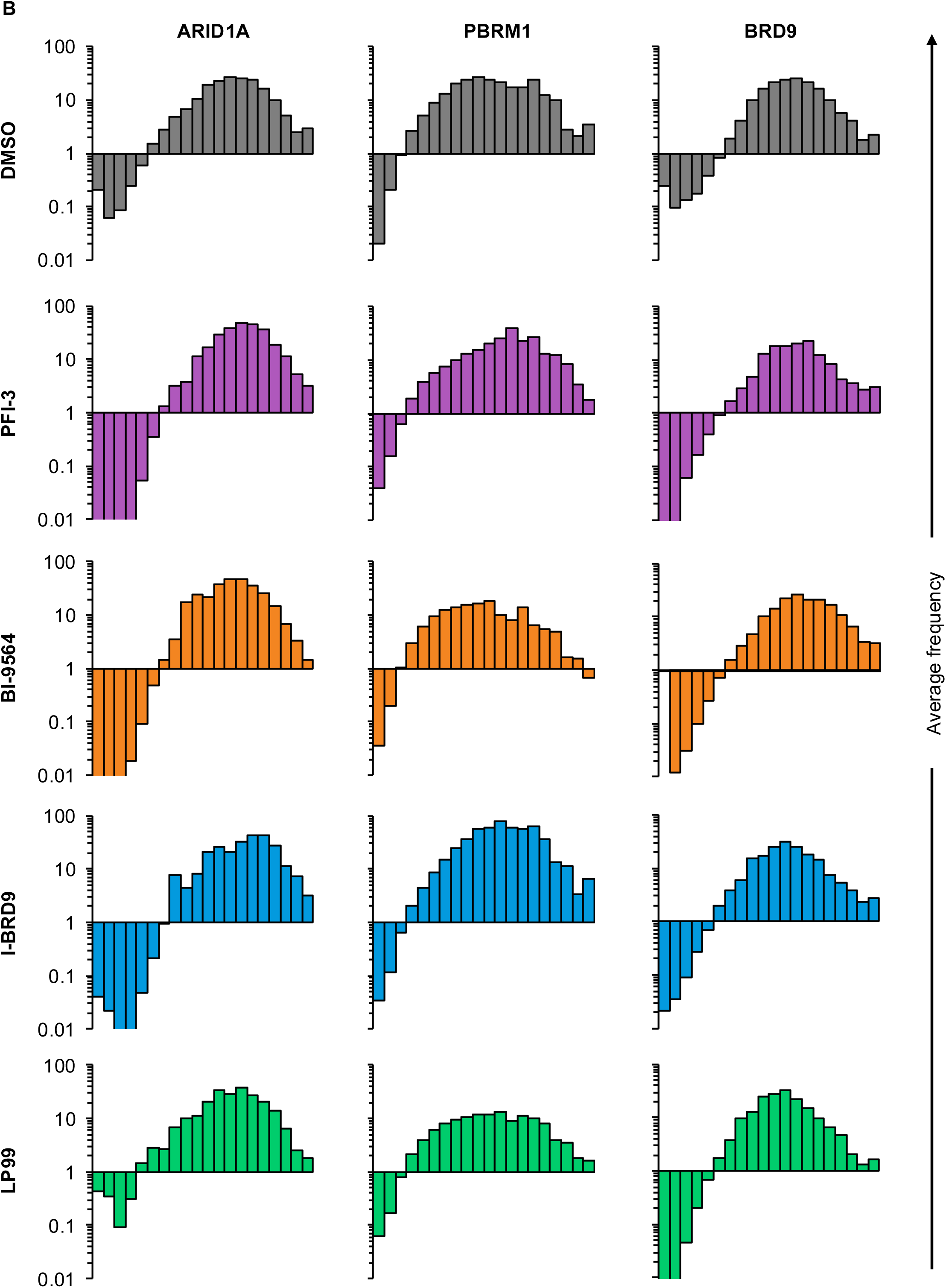
Distribution of unique BAF subunits during HSV-1 infection in the presence of small molecule bromodomain inhibitors. **A.**Representative micrographs of HeLa cells infected with HSV-1, treated with DMSO vehicle, LP99 (10 μM), PFI-3 (32 μM), BI-9564 (15 μM), or I-BRD9 (10 μM), fixed at 6 hpi, and stained for DAPI, HSV-1 ICP4, and the subunit of interest; n=2. **B.** Histograms presenting the average signal distribution of the unique BAF subunits in the presence of each inhibitor; n=2.

**Figure 10.**
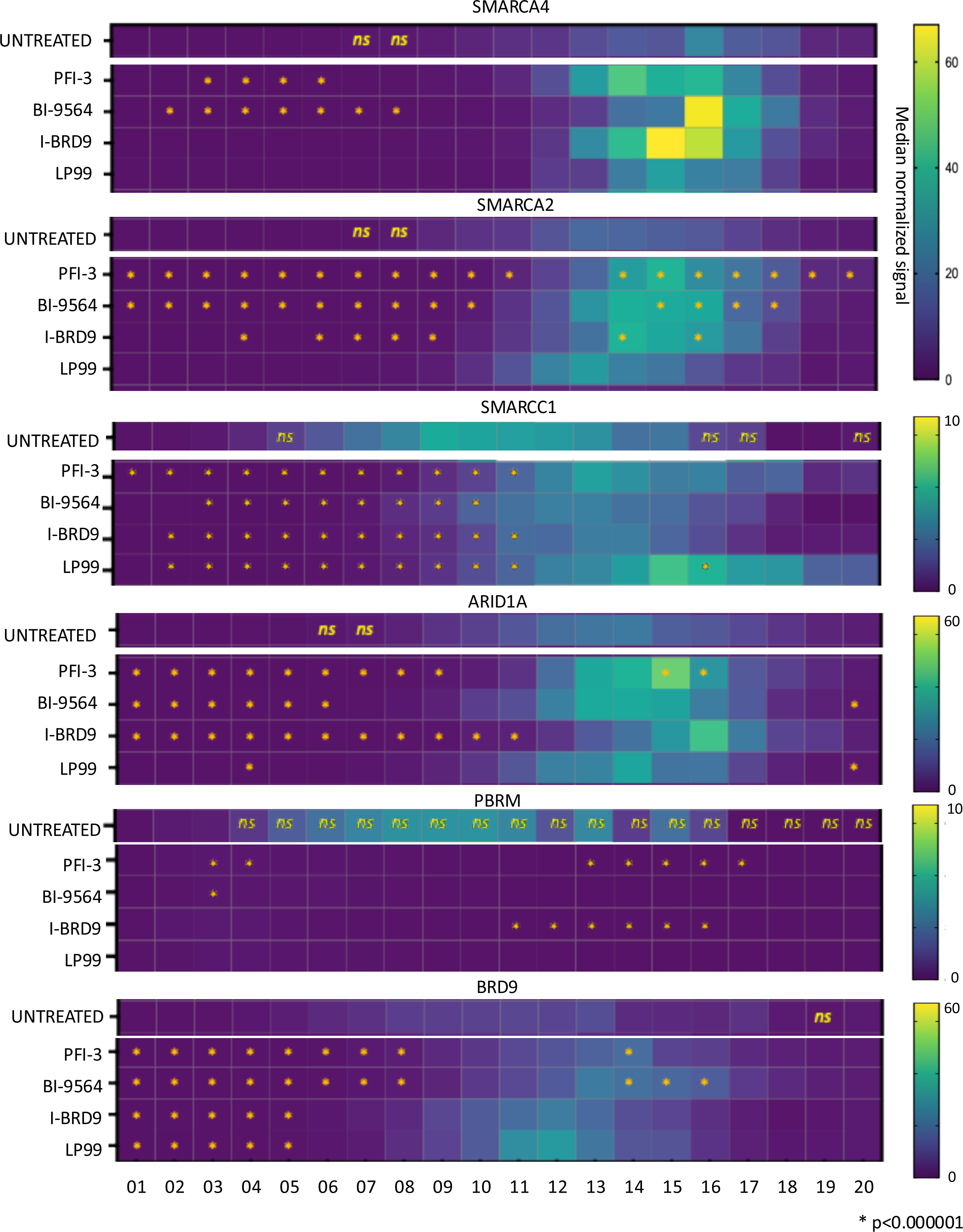
Small molecule bromodomain inhibitors do not deplete BAF subunits from the HND during HSV-1 infection. Heat map presenting the median normalized signal intensity in the different bins for the unique and shared BAF subunits in the presence of each inhibitor, in comparison to the distribution in untreated cells (copied from figure 4 for reference). *ns,* not significantly different from even distribution (untreated; Mann-Whitney, p >0.05); *, statistically different from the distribution in the untreated cells (treated cells; Mann-Whitney, p <0.05); n=2.

**Table 4.** Small molecule bromodomain inhibitors do not affect total subunit signal of shared BAF subunits within HND. Analysis of IF experiments described in figure 11. Total signal within HND regions divided by total signal within nuclear, non-HND regions, normalized by HND and nuclear, non-HND sizes. >1, overrepresentation of signal in HND. average ± standard deviation

| Treatment | SMARCA4 |  |  | SMARCA2 |  |  | SMARCC1 |  |  |
| --- | --- | --- | --- | --- | --- | --- | --- | --- | --- |
|  | AVG | SD | n | AVG | SD | n | AVG | SD | n |
| DMSO | 1.43 | 0.09 | 3 | 1.4 | 0.1 | 3 | 1.31 | 0.11 | 3 |
| LP99 | 1.38 | 0.05 | 2 | 1.33 | 0.16 | 2 | 1.4 | 0.01 | 2 |
| PFI-3 | 1.38 | 0.01 | 2 | 1.46 | 0 | 2 | 1.3 | 0.05 | 2 |
| BI-9564 | 1.46 | 0.07 | 2 | 1.32 | 0.03 | 2 | 1.28 | 0.04 | 2 |
| I-BRD9 | 1.38 | 0 | 2 | 1.35 | 0.05 | 2 | 1.28 | 0 | 2 |

**Table 5.** Small molecule bromodomain inhibitors do not affect total subunit signal of unique BAF subunits within HND. Analysis of IF experiments described in figure 11. Total signal within HND regions divided by total signal within nuclear, non-HND regions, normalized by HND and nuclear, non-HND sizes. >1, overrepresentation of signal in HND. average ± standard deviation.

| Treatment | ARID1A |  |  | PBRM1 |  |  | BRD9 |  |  |
| --- | --- | --- | --- | --- | --- | --- | --- | --- | --- |
|  | AVG | SD | n | AVG | SD | n | AVG | SD | n |
| DMSO | 1.46 | 0.12 | 3 | 1.43 | 0.11 | 3 | 1.39 | 0.16 | 3 |
| LP99 | 1.36 | 0.22 | 2 | 1.47 | 0.08 | 3 | 1.39 | 0.04 | 2 |
| PFI-3 | 1.42 | 0.06 | 2 | 1.46 | 0.11 | 3 | 1.3 | 0.06 | 2 |
| BI-9564 | 1.36 | 0.03 | 2 | 1.47 | 0.07 | 3 | 1.39 | 0.25 | 2 |
| I-BRD9 | 1.46 | 0.15 | 2 | 1.63 | 0.27 | 3 | 1.26 | 0.23 | 2 |

### BAF bromodomain are involved in inhibiting HSV-1 transcription but not DNA replication

The bromodomains of BAF subunits also interact with transcription activators and proteins involved in DNA replication and coordinate transcription to DNA repair (36, 47–49). Having established that the BAF complexes are not recruited to the HND by their bromodomains interacting with acetylated histones and knowing that they interact with two HSV-1 proteins involved in activation of transcription and DNA replication, we next assessed their effects on IE, E, E/L, and L transcription, and DNA replication, as well as viral assembly and egress. The expectation was that complexes that are recruited to the HND by viral proteins and regulate transcription and DNA replication and would likely enhance viral transcription or DNA replication.

RNA was isolated at 3, 6, and 9 hpi and mRNA levels were assessed by reverse transcriptase-quantitative PCR (RT-qPCR). Contrary to expectations, all small molecule bromodomain inhibitors increased RL2 mRNA (IE) abundance over the DMSO-treated controls at 3 6, and 9 hpi (p <0.01, 0.05, or 0.01, respectively, figure 11). However, the size effect at 9 hpi was minimal at about only 1.3-fold increase in mRNA levels (figure 11). Treatment also increased UL23 mRNA (E) at 6 or 9 (p <0.05), but not at 3 hpi. The effect was less marked than for the IE transcript, at about only 2-fold increase. E/L transcription does not require DNA replication but increases with HSV-1 genome copy number. mRNA levels of the E/L gene UL46 increased in infected cells treated with the inhibitors at 3, 6, and 9 hpi (p= 0.01, 0.05, or 0.05, respectively - figure 11).

**Figure 11.**
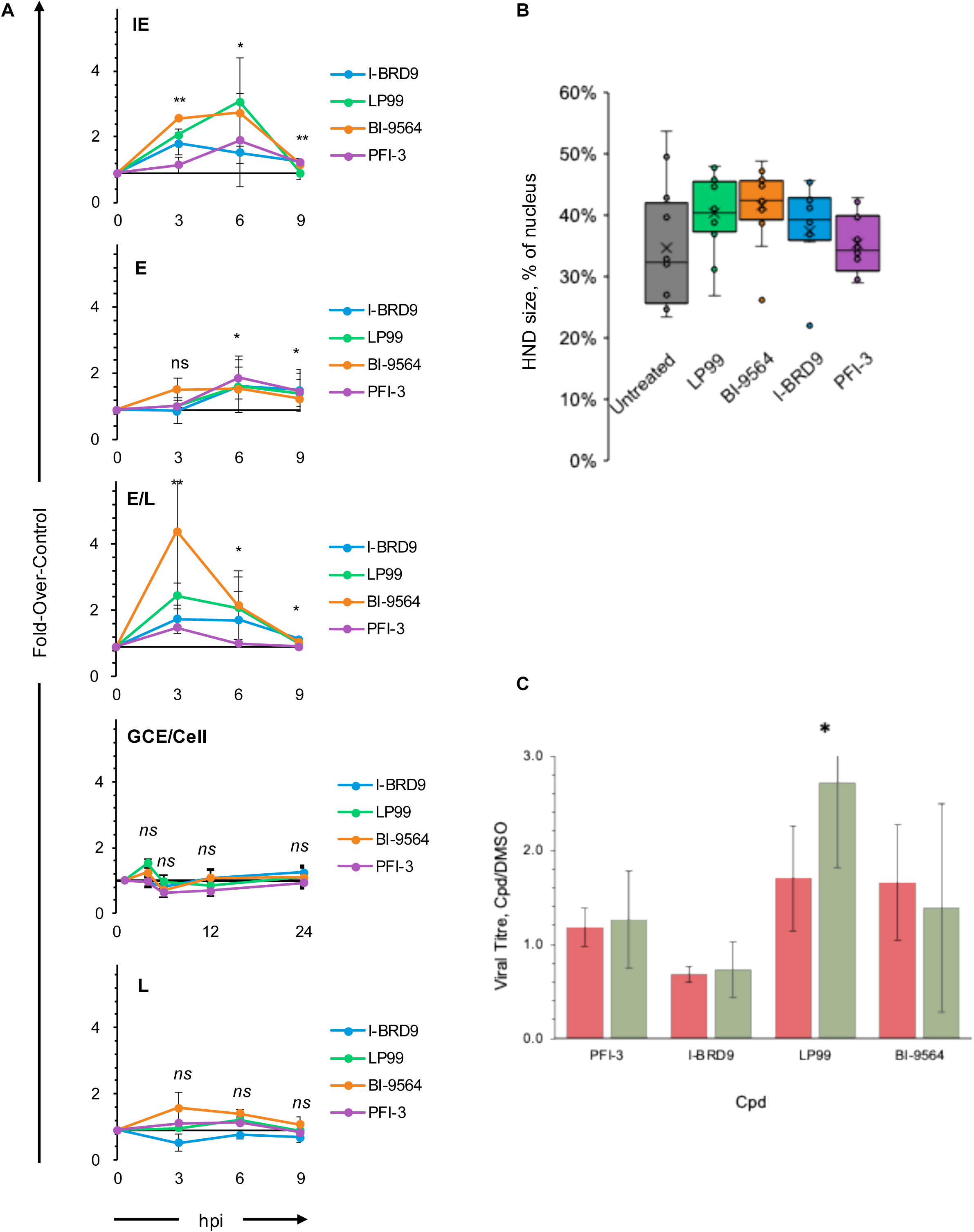
Small molecule bromodomain inhibitors modulate transcription of immediate early, early, and early late genes but not true late genes or DNA replication. **A.**HeLa cells were infected with HSV-1 KOS (MOI 5) and treated with DMSO vehicle, LP99 (10 μM), PFI-3 (32 μM), BI-9564 (15 μM), or I-BRD9 (10 μM). RNA was isolated at 3, 6, and 9 hpi. DNA was isolated at 2, 4, 6, 12, and 24 hpi. Relative mRNA abundance or genome copy number was assessed through quantitative PCR, normalized to the DMSO vehicle treated infections at each time point. Immediate early, RL2/ICP0. Early, UL23/TK. Early late, UL46/VP11. Late, UL44/gC. n=3, average ±SEM. **B.** Size of the HND at 6 hpi. The size of the HND in the cells analyzed in Figures 9 and 10 is plotted as box and whiskers against treatment. There were no statistically significant differences across the treatment groups. **C.** Viral burst assay at 18 hpi in HeLa (green) or HFF (red) cells treated with the different compounds normalized to the DMSO vehicle treated controls; n=2-4 for each compound and cell, average ±SEM. *, p<0.05.

Early proteins are required for DNA replication. We thus tested whether the increases in E mRNAs led to increases in viral genome copy number. DNA was isolated at 2, 4, 6, 12, and 24 hpi, and genome copy number was evaluated by quantitative PCR (qPCR) . The inhibitors did not affect viral genome copy equivalent (GCE) number per cell at any time, as shown by the lack of difference over the DMSO controls (p >0.05, figure 10). DNA replication is required for transcription of true L genes. As expected from their lack of effect on DNA replication, the inhibitors did not affect the levels of the L transcript evaluated (UL44) either.

There were statistically significant differences on the size effects of each inhibitor on IE, E/L and L mRNA levels (p<0.05, ANOVA), but not on the E one tested (p=0.13, ANOVA). The effects of LP99 and BI-9564 on all genes correlated the most, and of LP99 with I-BRD9 the second most (r^2^ 0.70 and 0.77, respectively), which is not surprising as LP99, BI-9564, and I-BRD9 target the bromodomains of BRD-9 and BRD-7 whereas PFI-3 does not. The effects of LP99 and BI-9564 on IE and E genes were highly correlated (r^2^ 0.75 and 0.84, respectively). For the E transcript, the effects of all four inhibitors were highly correlated, with the lowest correlation between BI-9564 and PFI-3 (r^2^ > 0.82 for all pairs except the later; 0.68 for the later). For the E/L mRNA, the correlation was high between LP99 and BI-9564 (r^2^ = 0.87) and I-BRD9 (r^2^ = 0.78).

Considering the changes in transcription of IE, E, and E/L genes, we evaluated the size of the HND in the cells infected with HSV-1 and treated with the different inhibitors which had been used for the analyses of the BAF subunit localization. Consistently with the effects on transcription, LP99 and BI9564 treatment resulted in the larger HND, whereas treatment with PFI-3 resulted in HND of similar sizes as in the vehicle treated controls (figure 11B). When considering the large distribution of HND size in the control cells, though, it is not surprising that the differences were not statistically significant (p>0.05).

As the inhibitors did not have a major effect on DNA genome copy number, or L gene transcription, they would not be expected to have a large effect on production of infectious virions. We nonetheless evaluated for any effects on viral assembly or egress by harvesting 18 hpi supernatant from HeLa or HFF cells infected with HSV-1 and treated with DMSO, PFI-3, LP99, BI-9564, or I-BRD9 (figure 11C). There was a mild increased viral production for LP99 and BI-9564, which reached (LP99) or approached (BI-9564) statistical significance at p<0.05, but not for PFI-3 or I-BRD9. The minimal changes in the release of infectious virions for one, or perhaps two, inhibitors only, and in the absence of increased viral genome copies, or L mRNA levels, suggests a likely indirect effect through modulation of cellular, not viral functions.

## Discussion

Here we show that the shared BAF ATPases SMARCA2 and SMARCA4 and scaffold SMARCC1, as well as a unique subunit from each cBAF, PBAF, and GBAF are enriched in the HND during lytic infection, mostly independently of any interactions between their bromodomains and acetylated histones. Together with previous results showing that some of the subunits directly interact with VP16 or ICP8, these results lead to the conclusion that the BAF complexes are recruited by viral proteins that activate viral transcription and DNA replication. Yet, they inhibit IE, E, and E/L mRNA accumulation. These effects are most consistent with the BAF complexes being recruited by the virus to inhibit widespread viral transcription through their interactions with transcription regulators at early times and then being counteracted by IE or E proteins or DNA replication. By evaluating the effects of the different inhibitors comparatively, GBAF or PBAF appear to be the BAF complexes most likely to be involved in the inhibition of viral transcription, which is consistent with their known activity inhibiting transcription at DNA damage sites in the cellular genome (36), the nicks and gaps in the HSV-1 genomes (50), and the induction of a DNA damage response by the infecting genomes (72). Considering that some subunits are recruited by the DNA replication protein ICP8, and BAF complexes inhibit transcription till DNA replication resumes, we favor DNA replication as the main counteractor of the inhibition of transcription by the BAF complexes. Inhibition of transcription by the BAF complexes, predominantly PBAF or GBAF, is thus a likely contributor to the regulation of the cascade of HSV-1 gene expression.

Taylor and Knipe showed that the shared BAF complex subunits SMARCA4/BRG1, SMARCA2/BRM, SMARCC1/BAF155, SMARCC2/BAF170, SMARCE1/BAF57, and SMARCD1/ BAF60a coprecipitate with the ssDNA-binding protein ICP8 during lytic infection in HEp-2 cells (51). Localization to replication compartments (a subset of HND) was shown for SMARCA4 and SMARCA2 (51). Herrera and Triezenberg reported that SMARCA4 and SMARCA2 bind IE loci during HSV-1 KOS infection. Binding was disrupted during infection with the HSV-1 VP16 activation domain (AD) mutant RP5, indicating that they are recruited through the VP16 AD (52). In a follow-up study, siRNA-mediated knockdown of SMARCA4 and SMARCA2 resulted in increased abundance of HSV-1 IE transcripts, even in the presence of cycloheximide (26), which is consistent with our results with small molecule inhibitors. Triezenberg and colleagues concluded that SMARCA2 and SMARCA4 are not required transcriptional coactivators of HSV-1, consistently with our findings. Their specific results are also consistent with ours showing that these BAF subunits are involved in repressing IE transcription. Differing from these early studies, we included specific subunits and inhibitors to evaluate cBAF, PBAF, and GBAF, complexes which had not yet been described when the previous works were published. We also used chemically unrelated small molecule bromodomain inhibitors to test the roles of their bromodomains. Differing from genetic intervention such as siRNA, small molecules allow inhibition of the targeted complexes only after infection and do not disrupt the complexes including the targeted protein, but rather disrupt their binding to acetylated residues. Also differing from previous work, we assessed the effects of the BAF complexes on IE, E, E/L, and true L mRNA accumulation, as well as genome copy number. We thus explored the roles of the different BAF complexes throughout infection, and consequently we reach the new conclusion that PBAF and GBAF complexes repress HSV-1 IE, E, and E/L transcription, but not DNA replication or L gene transcription. The use of inhibitors also allowed us to reach the new conclusion that their recruitment is independent of interactions between the bromodomains and acetylated histones and therefore depend on the previously reported interactions with ICP8 and VP16.

We used HeLa and human foreskin fibroblast (HFF) cells as complementary models because each of them recapitulates different aspects of natural infection. HeLa cells are transformed cervical epithelial cells, and HFF cells are primary human fibroblasts, thus providing histologically relevant and primary cells, respectively. The types of analyses performed here cannot be performed in stratified epithelial cells, and the consistency of results across a transformed epithelial cell line and human primary cells indicates that these results are not cell-type dependent.

Historically, the BAF ATPases have been regarded as interchangeable, although recent evidence suggests that they have unique roles in the cellular genome. SMARCA2 and SMARCA4 mutants have unique phenotypes, and their bromodomains show different binding preferences in vitro (53, 54). We found no differences in enrichment between SMARCA2 and 4 in either HeLa or HFF cells, suggesting that the two may be equivalent in their roles in HSV-1 replication.

The enrichment of SMARCA4 in HND, and the availability of a validated SMARCA4 antibody (40, 55), led us to test the binding of SMARCA4 to HSV-1 genomes using ChIP-qPCR. For comparison we included several human loci. Interleukin 24 (IL24) is a cytokine predominantly produced by T but not HeLa cells (56). Leucine rich repeat containing G protein-coupled receptor 6 (LGR6) is a ubiquitously expressed glycoprotein hormone receptor expressed at low levels (5.2 nTPM) in HeLa cells (43), as is the neuronal channel NPTXR. The chromosome 14 (chr14) non-coding locus is within an intergenic region between SERPINA1 and SERPINA11 and that in the chromosome 2 is in a gene desert. Glyceraldehyde-3-phosphate dehydrogenase (GAPDH) is a highly expressed (6,000 nTPM) housekeeping gene in HeLa cells (57). In a previous study, SMARCA4 bound to LGR6, IL24, and the chromosome 14 intergenic region at a rate of roughly one to five percent in NCI-H1944 adenocardinoma cells (40). Consistent with previous results (40), SMARCA4 bound to about 10% of each cellular loci (12.65% average), and likely even better to the gene desert in chromosome 2, although our studies were not powered to evaluate such differences. We found that HSV-1 infection and replication does not disrupt binding of SMARCA4 to its cellular loci. Although the difference in binding between cellular and viral loci did not reach statistical significance, it was consistently about three-fold lower for all viral than cellular loci, except for the gene desert in chromosome 2. Four million cells were analyzed in each ChIP, yielding at most 1.6 x 10^7^ potential binding sites for each cellular locus (assuming all cells in G2) and at least 5.2 x 10^9^ potential binding sites for each viral locus (assuming 1.3 x 10^3^ HSV-1 genomes per cell and single copy genes). Considering the co-IP efficiencies, SMARCA4 bound at most 1.6 x 10^6^ copies of each cellular site tested (i.e., less than one site per cell on average) and at least 2.6 x 10^8^ copies of each viral site tested. SMARCA4 thus bound on average 100-fold more of each viral than cellular locus. When considering the 10,000-fold difference in genome size, and assuming random distribution of binding sites, 99.99% of SMARCA4 in infected cells binds to cellular loci, and only 0.01% to viral ones. A significant fraction of each BAF complex is freely diffusing throughout the nucleus in uninfected cells (44, 46, 58), which may well be the complexes recruited by VP16 and ICP8 (24, 29), which are themselves enriched in the HND (28, 59, 60), with no need to deplete binding to the cellular genome. Our current analysis focused on specific cellular and viral loci but HSV-1 genome accessibility and chromatinization are regulated at the genome-wide level (7). We would thus expect BAF complexes to bind with equivalent efficiency at loci throughout the HSV-1 genome as well.

We used PFI-3, LP99, BI-9564, and I-BRD9. PFI-3 targets both mutually exclusive ATPases SMARCA2 and SMARCA4, which are incorporated into all BAF complexes, and PBRM1, which is only in PBAF. LP99, BI-9564, and I-BRD9 preferentially target PBAF and GBAF through BRD7 and BRD9, respectively, with some differences in potency against each of them (44, 46, 58). These inhibitors are structurally independent, target different bromodomains, and have been extensively validated. Their selectivity has been rigorously tested against bromodomain (BRD) families I-VIII in vitro (44, 46, 58, 61). In cultured cells, PFI-3 and BI-9564 increase the dynamics of their target subunits, while LP99 disrupts BRD7 and 9 interactions with histones (44, 46, 58, 62). It is unlikely, albeit not impossible, that the effects observed in this study are due to common off-target effects of the four different inhibitors, and it is also possible, even likely, that at least some of the observed effects are mediated by changes in cellular gene expression. Small molecule inhibitors have the advantage over genetic interventions in this respect in that they are added after infection, thus minimizing the time to act via such indirect effects.

PFI-3 targets two bromodomains in PBRM1 but did not affect its localization. PBRM1 encodes eight bromodomains, only two of which are targeted by PFI-3 (figure 7B). PBRM is a component of PBAF, which includes BRD7 with yet another bromodomain. GBAF and cBAF have just two and one bromodomains, respectively (figure 7B). These results would suggest that the non-targeted bromodomains in PBAF may compensate in part for the ones inhibited by PFI-3. Moreover, PBRM1 also binds to DNA directly, which is independent of any bromodomain. In any event, the changes in mRNA levels did not correlate with the changes in subunit localization. LP99 had the greatest influence on mRNA levels (figure 11), whereas BI-9564 had the largest effects on localization of most subunits (figure 10). As a caveat, some of the changes in mRNA levels were observed at 3 and 9 hpi, whereas localization was tested at 6 hpi. LP99 might affect localization at earlier (3 hpi) or late (9 hpi) but not intermediate (6 hpi) times, for example, whereas BI-9564 may have the largest effect at later times.

Histone acetylation inhibits formation of higher order chromatin structures and is subsequently associated with accessible and transcribed chromatin. HSV-1 chromatin during lytic infection is highly dynamic and associated with acetylated histones, including H3 lysine 9, 14, and 27 (7, 10, 30, 52, 63–67). Histone acetylation within the viral genome, however, does not recapitulate that within the cellular genome, in which it is largely limited to active promoters and enhancers; instead, histone acetylation is distributed throughout HSV-1 genomes, regardless of transcriptional state of individual loci (reviewed in (7, 9)). Total histone acetylation of the viral chromatin increases as infection progresses and chromatin dynamics increase (63). Trimethylation of H3K9 and H3K27, which are typically associated with heterochromatin, also increase in HSV-1 genomes as infection progresses (63). We characterized the localization of four major histone H3 acetylation marks: K4, K9, K27, and K36, normalized to total H3. Although acetylated histones promote chromatin dynamics and transcription within the cellular genome, associate with HSV-1 genomes, and contribute to BAF complex binding and stabilization, these four major histone acetylation marks were relatively depleted from HND with regards to total H3, suggesting that they are not a major contributor to the recruitment of BAF complexes to HND. Other PTM, such as H3K56ac (68, 69), may also recruit BAF complexes and BRD7 interacts, when not in BAF complexes, with H4K16ac (70), which may also direct its recruitment. We disfavor this possibility because of the known interactions of BAF subunits with viral proteins that localize to the HND. Moreover, four inhibitors that are specific for given bromodomains but not for the acetylated histone residues they bind to did not impede the recruitment of the subunits to the HND either.

Our findings that histone acetylation marks known to recruit BAF complexes are relatively depleted from HND and that bromodomain inhibitors do not decrease BAF subunit localization to HND, along with the previous work by other groups showing that BAF complexes are recruited by the VP16 AD and interact with ICP8, indicate that BAF complexes are recruited to HND through viral proteins that activate transcription or DNA replication. Their inhibitory activity on HSV-1 transcription is consistent with a recently reported similar inhibition of induced host gene transcription by BAF (47), and their well-established inhibition of transcription around sites of DNA damage (36, 49). The infecting HSV-1 DNA has nicks and gaps (71), and infecting HSV-1 DNA activates the DNA damage response (72), which is consistent with their ability to inhibit HSV-1 DNA transcription. It may still appear surprising that these complexes are recruited by viral proteins to silence the viral genome at IE and E times. However, viruses often suppress their own transcription to achieve proper replication kinetics. For example, HSV-1 ICP22 limits RNA polymerase II processivity to limit cryptic and anti-sense transcription (73).

Although HSV-1 genomes are chromatinized and associated with epigenetic marks including histone variants and post-translational modifications, HSV-1 transcriptional regulation is markedly different from that of the cellular genome. Epigenetic modifications such as histone variants and post-translational modifications appear to regulate HSV-1 transcription at the genome-wide level, while the individual promoters regulate the transcription of each gene. A major outstanding question about the regulation of HSV-1 gene expression is how the onset of DNA synthesis licenses the transcription of true L genes. Transcription of IE, E, and E/L mRNA were dependent on BAF complexes, but DNA replication or true L mRNA levels were not. BAF inhibition of HSV-1 transcription may well be limited to prior to the onset of DNA synthesis because the disruption of the HSV-1 DNA nucleoprotein complexes during DNA replication prevents further inhibition by chromatin.

We propose a model in which VP16 and ICP8 recruit BAF complexes to HND to inhibit most viral transcription, by interacting with transcription factors via their bromodomains, before the onset of DNA replication to help regulate the kinetics of gene expression (24, 26, 29, 52, 74). DNA replication releases this inhibition, and the BAF complexes then have no effect on true L gene expression. The BAF complexes therefore are some of the factors that need to be titered out by DNA replication to allow expression of late genes.

The development of small molecule bromodomain inhibitors has largely been driven by the high prevalence of mutations in chromatin remodeling complexes in cancers (19, 21, 40). Based on the results presented here, it is important to consider that the use of such anti-cancer drugs may enhance HSV-1 or -2 gene expression, or perhaps even induce reactivation from latency; pre-emptive antiherpetic therapy may be considered to minimize this possibility.

## Acknowledgements.

This work was funded by the NIAID of the NIH (1R01AI153396) to LMS. SMS work was partly supported by a scholarship from the Center for Vertebrate Genomics (CVG), Cornell University. LMS wishes to acknowledge the generous support from an anonymous donor.

## Materials and Methods

### Cells

HeLa cells were obtained from ATCC (CCL-2). Human foreskin fibroblast (HFF) cells were obtained from ATCC (SCRC-1041). African green monkey kidney (Vero-76) cells were obtained from ATCC (CRL-1587). HeLa, HFF, and Vero-76 cells were maintained in 5% fetal bovine serum (FBS) in Dulbecco’s modified Eagle medium (DMEM).

### Virus

Wild-type herpes simplex virus 1 (HSV-1) KOS strain was obtained from the late Dr. P. Schaffer, University of Pennsylvania, Philadelphia, PA, US and has been previously described(50). Viral stocks were maintained and titrated on Vero-76 cells.

### Compounds

50 mM stocks of I-BRD9, PFI-3, LP99, and NVP-RXI570 were prepared in DMSO. 15 mM stocks of BI-9564 were prepared in DMSO. Each aliquot was stored at −20°C and thawed no more than 4 times. Catalog numbers and working concentrations are listed in table 5.

### Immunofluorescence

HeLa or HFF cells were seeded onto round 18 mm high tolerance glass coverslips (Warner Instruments CS-18R17) in a 12-well plate (1 ml/well) at a density of 1.8 x 10^5^ cells/well or 0.9 x 10^5^ cells/well, respectively. After allowing cells to attach, cells were infected with 100 μl/well HSV-1 KOS inoculum prepared at a multiplicity of infection (MOI) of 5. Plates were rocked and rotated every 10 minutes for an hour, then the inoculum was washed away with 2 ml/well cold serum-free media (SFM) and replaced with 1 ml/well warm 5% FBS DMEM. For studies using small molecule bromodomain inhibitors, cells were treated with 30 μM PFI-3, 10 μM LP99, 15 μM BI-9564, 10 μM I-BRD9, or 0.1% DMSO in 5% FBS DMEM.

Cells were fixed at 6 h post infection (hpi) with 2 ml/well 4% formaldehyde 4% sucrose in phosphate-buffered saline (PBS; Corning 21-040-CV) for 10 minutes at 4°C with gentle rocking. Cells were washed thrice in 0.1% BSA in PBS for 5 minutes each at room temperature with gentle rocking, then permeabilized with 1 ml/well 0.2% Triton X-100 in PBS for 10 minutes at 4°C with gentle rocking. Cells were washed twice in 0.1% BSA in PBS for 5 minutes each at room temperature with gentle rocking, then blocked with 500 μl/well 0.1% Triton X-100 1% BSA 5% normal goat serum (NGS) in PBS for 2 hours at room temperature with gentle rocking. Blocking buffer was removed and cells were incubated with primary antibody diluted in 350 μl/well 0.1% BSA 1% NGS in PBS overnight at 4°C with gentle rocking. Table 6 lists the primary antibodies used. Cells were washed thrice in 0.1% BSA in PBS, then incubated with secondary antibody in 0.1% BSA 1% NGS in PBS at room temperature with gentle rocking. Coverslips were protected from light following the addition of secondary antibodies. Table 6 lists the secondary antibodies used. After 50 minutes, 1 μg/ml DAPI for 10 minutes was added. After 10 minutes, secondary antibody and DAPI solution was removed, and coverslips were washed twice with 0.1% BSA in PBS and once with water. Coverslips were mounted with ProLong (ThermoFisher P36930). Images were taken on an Olympus Confocal microscope at 60X and processed using FIJI. At least eight, and typically more, randomly selected nuclei were analyzed from each biological replicate in each cell line, for a total of at least 32 nuclei per treatment, and typically more. The number of nuclei analyzed was based on the highly reproducibility across nuclei, cell lines, and experiments, and as shown by the statistical power to identify differences among proteins. The differences observed across different proteins indicate that any potential signal bleed through from the ICP4 antibody does not significantly influence the outcomes.

**Table 6.** Small molecule bromodomain inhibitors used in this study. Working concentration refers to range of concentrations used for the viral burst assay (figure 9); highest working concentration used for all other experiments.

| Compound | Source | Catalog No. | [Stock] | [Working] |
| --- | --- | --- | --- | --- |
| LP99 | Cayman | 17661 | 50 mM | 0.3 $\mu\text{M}$ – 10 $\mu\text{M}$ |
| PFI-3 | Selleck | S7315 | 50 mM | 0.3 $\mu\text{M}$ – 32 $\mu\text{M}$ |
| BI-9564 | Selleck | S8113 | 15 mM | 0.3 $\mu\text{M}$ – 15 $\mu\text{M}$ |
| I-BRD9 | Selleck | S7835 | 50 mM | 0.3 $\mu\text{M}$ – 10 $\mu\text{M}$ |

**Table 7.** Antibodies used in this study.

| Antigen | Antibody | Source | Host | Dilution |
| --- | --- | --- | --- | --- |
| HSV-1 ICP4 | ab6514 | Abcam | Mouse | 400 |
| SMARCA6 | ab300040 | Abcam | Rabbit | 1200 |
| SMARCA2 | ab240648 | Abcam | Rabbit | 1200 |
| SMARCA4 | ab110641 | Abcam | Rabbit | 1200 |
| SMARCC1 | ab172638 | Abcam | Rabbit | 1200 |
| ARID1A | ab182560 | Abcam | Rabbit | 1200 |
| PBRM1 | ab243876 | Abcam | Rabbit | 600 |
| BRD9 | AB_2856283 | Invitrogen | Rabbit | 600 |
| H3K4ac | ab176799 | Abcam | Rabbit | 2400 |
| H3K9ac | ab32129 | Abcam | Rabbit | 1200 |
| H3K27ac | ab4729 | Abcam | Rabbit | 2400 |
| H3K36ac | ab177179 | Abcam | Rabbit | 1200 |
| Total H3 | AB_2687567 | Invitrogen | Rat | 600 |
| Ⓜ-Rabbit Alexa Fluor 488 | AB_143165 | Thermo Fisher | Goat | 2000 |
| Ⓜ-Mouse Alexa Fluor 546 | AB_2534071 | Thermo Fisher | Goat | 2000 |
| Ⓜ-Rat Alexa Fluor 647 | AB_141778 | Thermo Fisher | Goat | 2000 |

### Image Analysis

Using Python, text image files of ICP4 and DAPI channels were used to mask herpes nuclear domains (HND) and nuclear non-HND regions, respectively. Values from the channel of interest were then sorted as inside or outside viral regions or discarded if extranuclear. R was used to plot histograms of frequency of pixel intensities, divided into 20 bins for each nucleus analyzed. For each bin, the viral region frequency was normalized by the host region frequency. For histone posttranslational modifications (PTM), this analysis was completed in each nucleus for the specific PTM and total histone H3. For each bin, the PTM ratio was normalized by the total histone H3 ratio.

### Chromatin Immunoprecipitation

Chromatin immunoprecipitation was performed as previously described with the following modifications (6). Sonicated chromatin was thawed on ice and brought to a final concentration of 833 ng/mL with ChIP binding buffer (1% Triton X-100, 10 mM Tris pH 8.0, 150 mM NaCl, 2 mM EDTA pH 8.0) to achieve 499.8 ng of chromatin per IP reaction. Per IP reaction, 1.9 μg SMARCA4 (Abcam, ab110641) or non-specific rabbit IgG (Abcam, ab171870) per ChIP reaction was conjugated to 18.75 μl protein A Dynabeads (Invitrogen, 10001D) for 2 hours at room temperature. Chromatin was incubated with bead-antibody complex overnight at 4°C with constant rotation. Isolated DNA was subject to qPCR with cellular and viral primers listed below.

### Cell Viability Assay

HeLa and HFF cells were seeded onto 96-well plates at a density of 1.1 x 10^4^ cells/well or 7.8 x 10^3^ cells/well, respectively (0.125 ml/well). After allowing cells to attach, cells were treated with each small molecule bromodomain inhibitor in quadruplicate. The CellTiter-Glo Kit (Promega G7570) was used to assess ATP levels at 0 and 18 hours post treatment.

### RNA Extraction and cDNA Synthesis

HeLa cells were seeded at a density of 3.6 x 10^5^ cells/well onto 6-well plates (2 ml/well). After allowing cells to attach for 4 h, cells were infected with 200 μl/well HSV-1 KOS inoculum at an MOI of 5. Plates were rocked and rotated every 10 minutes for an hour, then the inoculum was washed away with 4 ml/well cold DMEM and replaced with 2 ml/well warm 5% FBS DMEM supplemented with 0.1% DMSO solvent or a dilution of a small molecule bromodomain inhibitor in DMSO (table 1). Cells were harvested for RNA extraction at 1, 3, 6, or 9 hpi. Media was removed and cells were washed with 1 ml/well PBS. Cells were scraped and transferred to screwcap tubes. Cells were pelleted at 500 g for 5 minutes at room temperature. Supernatant was discarded and cells were flash-frozen in liquid nitrogen and stored at -80C. RNA was isolated using the RNeasy Kit (Qiagen 74104) and DNA was degraded using the RNase-Free DNA Set (Qiagen 79254) following the manufacturer’s protocol. RNA was eluted in 30.0 μl H_2_O and stored at -80°C. The iScript cDNA Synthesis Kit (BioRad 1708890) was used to generate cDNA following the manufacturer’s recommendations using 2.0 µl RNA per reaction. Reverse transcription reactions were run with and without reverse transcriptase to control for any DNA recovered and not degraded during the RNA extraction.

### DNA Extraction

HeLa cells were seeded at a density of 3.6 x 10^5^ cells/well onto 6-well plates (2 ml/well). After allowing cells to attach for 4 h, cells were infected with 200 μl/well HSV-1 KOS inoculum at an MOI of 5. Plates were rocked and rotated every 10 minutes for an hour, then the inoculum was washed away with 4 ml/well cold DMEM and replaced with 2 ml/well warm 5% FBS DMEM supplemented with 0.1% DMSO solvent or a dilution of a small molecule bromodomain inhibitor in DMSO (table 1). Cells were harvested for DNA extraction at 1, 4, 6, 12, or 24 hpi. Media was removed and cells were washed with 1 ml/well PBS. Cells were scraped and transferred to screwcap tubes. Cells were pelleted at 500 g for 5 minutes at room temperature. Supernatant was discarded and cells were flash-frozen in liquid nitrogen and stored at -80C. DNA was isolated using the DNeasy Blood and Tissue Kit from Qiagen (69504) following the manufacturer’s protocol. DNA was eluted in 100.0 μl H_2_O and stored at -80°C.

### Primers

GAPDH: TTGGCTACAGCAACAGGGTG; GGGGAGATTCAGTGTGGTGG

LGR6: TGGGCTATGTGACCTTAGGC; CCCCAGGTCAGTGAGTTCAT

IG Chr14: CTCCCCTGGCATATTACCAA; GACCATTGTTCCCTCCTCAA IL24: GCACGACTCCTGGGCCGTAACG; ACGGGAGTAGGAATGTGACG

NPTXR: TCTGTGTTCCGGGGACGAT; CCTGGAGACAAAGGGTG

GD Chr2: AGTCATGTCACAGCATGGGT; CCTGGCCAACTTTTTGCATT RL2: CCCACTATCAGGTACACCAGCTT; CTGCGCTGCGACACCTT

UL23: ACCCGCTTAACAGCGTCAACA; CCAAAGAGGTGCGGGAGTTT

UL46: TCCTCGTAGACACGCCCCCCGT; ACGCCCCCTACGAGGACGACGAGT

UL44: GTGACGTTTGCCTGGTTCCTGG; GCACGACTCCTGGGCCGTAACG

LAT: CCAGGCAGTAAGACCCAAGC; GGCCGGTGTCGCTGTAAC

